# Environmental drivers of low vaccine responsiveness in a lab-to-wild rodent model

**DOI:** 10.1101/2025.10.20.683371

**Authors:** Simon A. Babayan, Saudamini Venkatesan, Jessica L. Hall, Ewan Smith, Amy Sweeny, Amy B. Pedersen

## Abstract

Vaccination is the most effective way to prevent infectious diseases and safeguard public health. Yet, most new vaccines fail in late clinical trials, and even established ones often underperform in populations apart from those in which they were initially tested. This can lead to reduced vaccine responsiveness, breakthrough infections, and prevent or delay herd immunity. While the causes of vaccine hyporesponsiveness remain difficult to identify, quantify, and therefore address, numerous reports indicate a predominant role of environmental factors. This has notably been demonstrated by a reduction in the immunogenicity and efficacy of various vaccines when transitioning from urban to rural human populations. Here, we tested whether and, if so, how the environment can cause vaccine hyporesponsiveness. We hypothesised that if the leading causes of vaccine hyporesponsiveness were environmental, then the reduced efficacy would be exacerbated when individuals are under nutritional stress; specifically predicting that high quality diet supplementation would increase vaccine responsiveness. Finally, we predicted that parasitic helminth infection, which are more common in rural populations, would degrade vaccine responsiveness, e.g. due to their ability to suppress host immunity, and that anthelmintic treatment could rescue vaccine responsiveness in infected individuals. To test these hypotheses, we coupled lab and field experiments with structural causal modelling, and measured antibody responses of paired conspecific cohorts of laboratory-reared and wild wood mice (*Apodemus sylvaticus*) to a single or two doses of diphtheria toxoid vaccine formulated with alum; with and without diet supplementation. We found that vaccine-specific IgG1 antibodies were ∼47% lower in the wild wood mice compared to the laboratory reared population. We also demonstrate that, across both habitats (wild and lab), substantial variation in vaccine responsiveness was caused by diet. However, contrary to our predictions, this high quality dietary supplementation resulted in lower vaccine responsiveness. Further, once the effects of habitat, diet, and sex were adjusted for, increasing helminth infection burdens negatively affected vaccine-specific antibody production. Using counterfactual modelling, we show that targeting anthelmintic treatment at heavily infected individuals would have improved their vaccine responsiveness by 2 to 4-fold. Our results indicate that the wild environment and access to a high-quality diet play a dramatic role in shaping the immune system’s response to immunisation. Further, we show that laboratory settings, even when using a genetic diverse, non-traditional model, may systematically overestimate vaccine performance by failing to capture natural environmental variability. Our study provides a robust, causally explicit modelling approach to disentangle and quantify the complex factors that drive vaccine responsiveness in focal populations. This framework offers a pathway for predicting vaccine performance across diverse populations and identifying targeted interventions to enhance immunisation outcomes in real-world settings.

## INTRODUCTION

Unanticipated variability in individual innate and adaptive immune responses to vaccination can lead to a lack of protection to the targeted disease, continued pathogen transmission across the wider population, and an increased risk of the evolution of vaccine escape [1, 2, 3, 4, 5]. Both intrinsic and extrinsic factors may impact vaccine responsiveness (defined here as the ability to generate vaccine-specific antibodies), including genetics [6] and epigenetics [7], sex [8], reproductive status [9], prior infections [10, 11, 12, 13, 14], and diet [15, 16, 17, 18]. Importantly, studies have shown that environmental factors may also be important [19, 20, 21], indeed, recent work demonstrated that even temporary exposure to a natural environment can drive B cell maturation and enhance antibody production [22]. Such negative impacts of the environment on the immune system’s response to vaccination has stark implications for human health and public health policy, manifesting for example in the substantial drop in vaccine responsiveness and efficacy in low-income or rural populations relative to high-income or urban populations [23, 19]. Furthermore, helminth infections, which disproportionately affect individuals in rural and low-income populations [24, 25], can cause poor responsiveness to vaccines due to their ability to systemically suppress their host’s immune system [26, 27] although the results are mixed [11].

As a consequence of these challenges, only a small minority of vaccines successfully transition from the preclinical stages of development through phase III trials to reach the global market [28], and there are still no antihelminthic vaccines approved for humans. Furthermore, even the most successful vaccines can suffer substantial loss of efficacy, notably for people of different ancestry [29] or different habitats than the populations in which they are initially deployed [19]. For example, individuals of Tanzanian ancestry were reported to have higher inflammatory and Type 1 immune markers [30] and responses to measles vaccination [31] than those of European ancestry. In contrast, the expression of BCG- and tetanus-specific cytokine recall and antibody responses in rural populations was found to be markedly lower than in urban populations, while humoral responses to tetanus vaccination were higher [21, 32]. Similar patterns have been observed across multiple vaccines [19]. Taken together, these studies indicate that vaccine efficacy depends on characteristics of both the vaccine and the recipient population, and highlights substantial gaps in our ability to predict when vaccines might provide effective immunity, and importantly when they may not, across different individuals and populations. Indeed, given the complex processes that regulate protective immune responses, it is unclear how all these factors interact with each other and the environment, and which among them could be targeted to maximise or improve vaccination efficacy.

To address these shortcomings and to increase the generalisability of experimental models used for immunology and vaccinology, recent efforts have been made to integrate natural variability into immunological research [33, 34, 35, 36, 37]. These include exposing mice to more diverse, “dirtier” environments, littermates, and wild-derived microbiota, to stimulate the development of more mature, and thus more representative, immune systems and responses to infection. However, there is still uncertainty about how these different sources of heterogeneity interact to affect the immune system [38], and what consequences this will have for vaccine responsiveness and the translation potential of mouse models for novel vaccines. Thus, even with more realistic lab models, it is still unclear how environmental factors and past stressors will influence responses to vaccination.

Further, there are statistical impediments to identifying the drivers of poor vaccine effectiveness. Indeed, observational ecological studies have often relied on statistical approaches that obscure causal relationships [39]. Here, to clarify the effects of key environmental variables, such as habitat, diet, and parasite infection, on vaccine responsiveness, here we combined immunisation experiments in both laboratory and field settings with structural causal modelling. Specifically, we conducted a randomised controlled vaccination and dietary supplementation experiment in a coupled lab-to-wild wood mouse *Apodemus sylvaticus* system. By using the same vaccine in a genetically outbred species across diverse habitats (lab-to-wild), we can isolate the environmental effects of living in the wild on vaccine responsiveness. For this, we used a Diphtheria toxoid vaccine to ensure no prior exposure in the wild population could bias the vaccine-specific responses. Further, we have previously demonstrated in this lab-to-wild model that high-quality diet supplementation drives a sustained reduction in helminth (*Heligmosomoides polygyrus*) burdens and transmission, while also improving the efficacy of anthelmintic treatment [40]. We therefore sought to test the predictions that vaccine responses in wild mice (i) would be lower than in lab-reared wood mice, and (ii) higher in diet-supplemented than normal-diet animals. Additionally, given extensive reports of helminths’ ability to suppress host immunity [41], we predicted that (iii) parasitic helminth burdens would reduce vaccine responsiveness, and (iv) that anthelmintic treatment could recover vaccine responsiveness in infected individuals. To test this latter prediction, we employed counterfactual modelling to simulate the effects of population-wide anthelmintic intervention on vaccine efficacy. To quantify the causal effects of habitat, diet, sex, and parasite infection on vaccine responsiveness, as measured by vaccine-specific IgG1, and identify the pathways through which their effects are mediated, we constructed a structural causal model (SCM) [42]. This causal framework uses graphical models and structural equations to inform statistical adjustments [43, 44] required for eliminating or reducing confounding, such as common cause (misattribution of causal effects), sampling bias (non-representative test population), and transportability bias (lack of generalisability). Under this framework, we used Bayesian models to estimate the average direct and indirect causes of lower vaccine responsiveness when translating a vaccine from the laboratory to the wild, while determining how uncertainty and biological variability impact the causal structure. This approach revealed surprising environmental drivers of vaccine failure (low responsiveness) and identified specific interventions that could dramatically improve vaccine performance in natural populations. Our findings challenge fundamental assumptions about vaccine development and provide a roadmap for designing more effective vaccination strategies for real-world deployment.

## RESULTS

### Vaccination induced weaker antibody responses in the wild than in the laboratory

Wild and laboratory wood mice (*A. sylvaticus*) were given a high- or low-quality diet and then randomly assigned to be immunised with a diphtheria-alum vaccine (DTV) once (non-boosted) or twice (boosted) or given a adjuvant-only control. DTV-specific IgG1 serum concentrations were then measured by ELISA seven to 35 days post immunisation to determine vaccine responsiveness (Figure 1; see details in “Materials, Methods, and Models”).

**Fig. 1:**
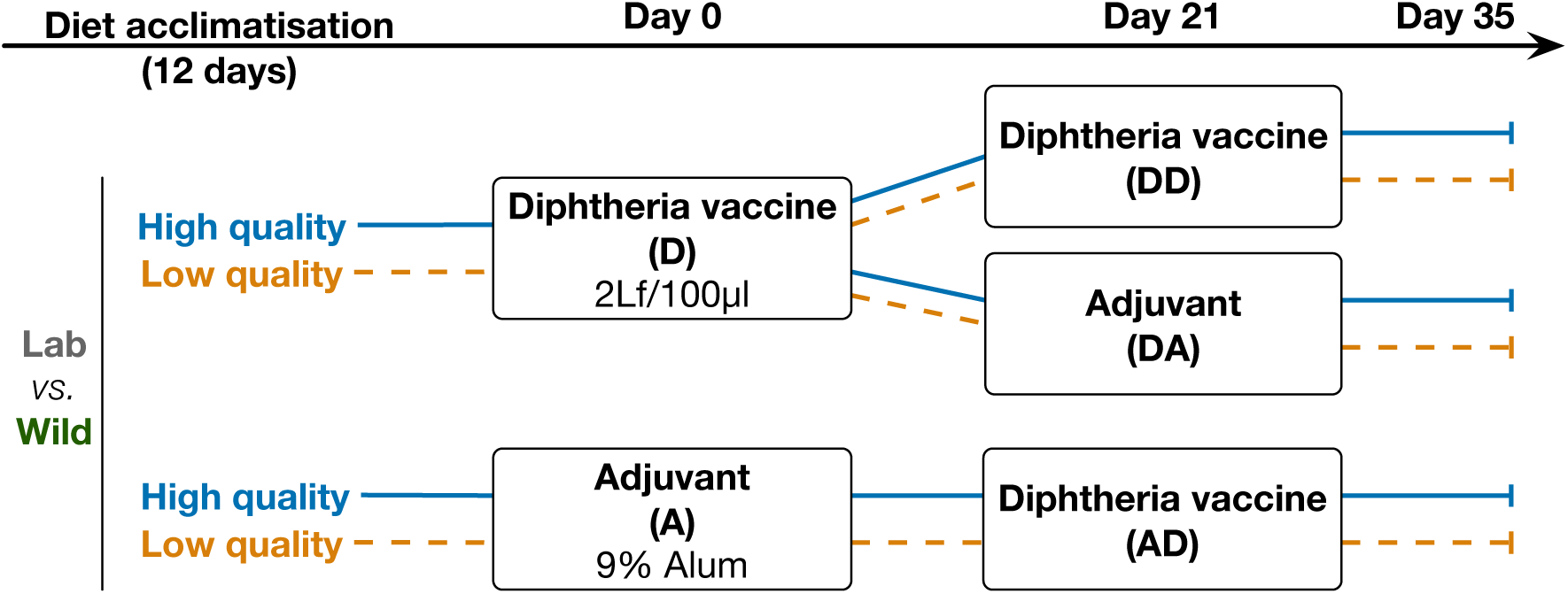
Experimental design. Wood mice from a wild-derived, now laboratory-reared, outbred colony and from the wild (Scottish woodlands) were provided enriched (TransBreed) or standard (lab: normal chow; wild: no supplementation) diets. After a minimum of 12 days, wood mice were allocated at random to immunisation with a 2Lf/100µl Diphtheria Toxoid + 9% alum vaccine (D) or with adjuvant-only control (A). Twenty-one days after their first immunisation, mice that had been immunised with their first dose (D) were then given a second dose (boost) of the vaccine (DD) or just given an adjuvant-only control (DA), while mice that had received an initial adjuvant-only control (A) were all given their first dose of 2Lf/100µl Diphtheria Toxoid + 9% alum vaccine for their second immunisation (AD).

DTV-specific IgG1 antibody responses were 46.9% ± 0.1% lower in the wild wood mice than their lab-based wood mouse conspecifics, regardless of the vaccine regimen given or diet (Figure 2A, 2B ‘Hab’), with sufficient variability to support strong conclusions about cause and effect (Supplementary Fig. S1A). While immunisation was most effective (as measured by responsiveness) with two doses across both lab and wild habitats, lab wood mice reached peak vaccine-specific antibody levels with only a single dose (Figure 2A, ‘DA’ vs ‘DD’; Lab boosted (two doses) vs. Lab non-boosted (one dose) OD = 0.2 ± 0.1, whereas Wild boosted vs. Wild non-boosted OD = 1.02 ± 0.23). However, in the wild, only mice given two doses (boosted) showed responsiveness similar to laboratory-housed wood mice (Figure 2A, ‘DD’; difference between Wild non-boosted vs. Lab non-boosted OD = 0.75 ± 0.11 while difference between Wild boosted and Lab boosted OD = 0.61 ± 0.18; LRT of Habitat ×vaccine regimen: Δ*dof* = 4, *χ*^2^ = 30.6, *p* < 0.001). Only wood mice receiving the diphtheria toxoid antigen generated DTV-specific IgG1, indicating no pre-existing immunity to this antigen in any of our study populations (Figure 2A, group ‘A’). Intriguingly, dietary supplementation appeared to reduce vaccine responsiveness in both habitats, though more strongly in the wild (Fig. 2B, Table 1).

**Fig. 2:**
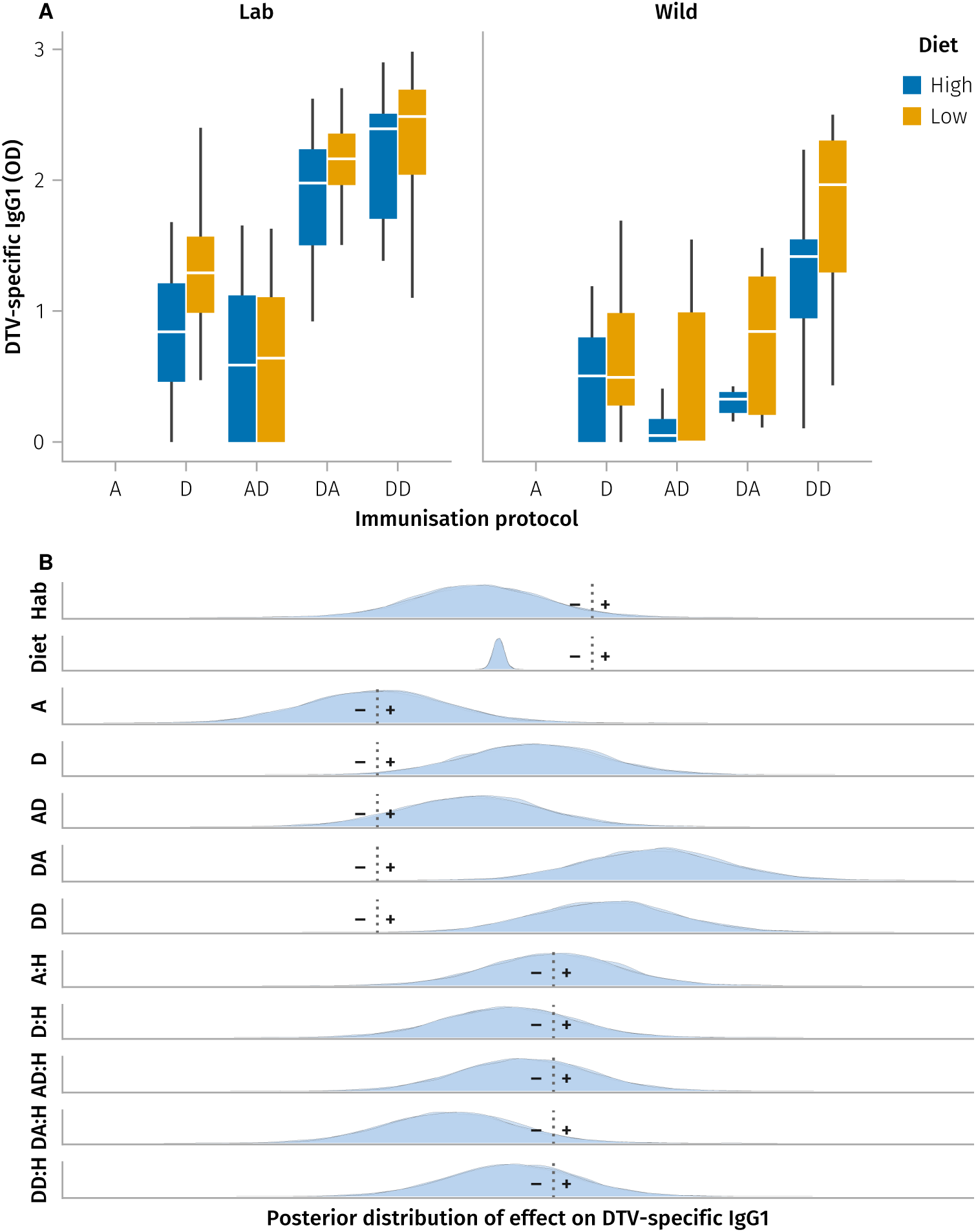
Correlates of vaccine responsiveness (DTV-specific IgG1) **A**, Diphtheria toxoid vaccine (DTV)-specific IgG1 optical density in wood mice from laboratory- and wild-habitats provided with either high or low quality/control diets. Mice were immunised either once, with adjuvant-only control (A) or DTV once (D), or twice, using adjuvant alone followed by DTV (AD), DTV and subsequently adjuvant alone (DA), or two separate doses of DTV (DD; boosted). Only serum samples taken >7 after immunisation are shown here. **B**, Posterior conditional distributions of the coefficients for Diet, habitat (Hab or H), and Vaccine formulation (A, D, AD, DA, DD), and interactions between each habitat and vaccine formulation (A:H, D:H, AD:H, DA:H, DD:H). Each density curve represents the distribution of 3,000 samples from one of four MCMC chains. The dotted line indicates the value for which the reference—population average vaccine-specific IgG1 values for Diet and Habitat, as well as average vaccine-specific IgG1 values for the Adjuvant alone control group—is zero (see parameter estimates in Table 1).

**Table 1:** To understand how different factors influence the vaccine-specific IgG1 immune response in individuals, we used a statistical model that accounts for individual differences and multiple categorical variables. The model assumes the immune response *E*_*i*_ for individual *i* follows a normal distribution with mean *α*_*ID*[*i*]_ *γ*_*H*[*i*]_ +*δ*_*D*[*i*]_ +*η*_*V*[*i*]_ +*θ*_(*V*×*H*)[*i*]_ and standard deviation *σ*. Here, *α*_*ID*[*i*]_ is the individual-specific baseline (random intercept), *γ*_*H*[*i*]_, *δ*_*D*[*i*]_, and *η*_*V*[*i*]_ are the fixed effects of habitat *H*, diet *D*, and vaccine formulation *V*, respectively, and *θ*_(*V*×*H*)[*i*]_ represents the interaction effect between vaccine and habitat. The model captures how these factors and their interaction contribute to variation in immune response.

| Parameter | Mean | Std | 5% | 50% | 95% |
| --- | --- | --- | --- | --- | --- |
| **Intercept** | 1.02 | 0.85 | -0.39 | 1.032 | 2.43 |
| **Habitat** | -0.519 | 0.806 | -1.863 | -0.513 | 0.795 |
| **Diet** | -0.253 | 0.072 | -0.373 | -0.253 | -0.137 |
| **A** | 0.0 | 0.871 | -3.307 | -1.892 | -0.438 |
| **D** | 2.062 | 0.856 | -1.228 | 0.175 | 1.598 |
| **AD** | 1.256 | 0.875 | -2.061 | -0.651 | 0.84 |
| **DA** | 3.652 | 0.87 | 0.343 | 1.751 | 3.215 |
| **DD** | 2.996 | 0.865 | -0.304 | 1.091 | 2.534 |
| **A:H** | 0.0 | 0.818 | -0.864 | 0.47 | 1.831 |
| **D:H** | -0.594 | 0.811 | -1.434 | -0.125 | 1.223 |
| **AD:H** | -0.371 | 0.823 | -1.244 | 0.103 | 1.472 |
| **DA:H** | -1.348 | 0.818 | -2.228 | -0.874 | 0.496 |
| **DD:H** | -0.527 | 0.818 | -1.389 | -0.061 | 1.31 |

To further quantify how habitat (lab *vs.* wild), diet (supplemented *vs.* control), and the immunisation protocol (non-boosted *vs.* boosted) affect variation in vaccine respons-iveness, we constructed a multilevel Bayesian model with varying intercepts for each immunisation regimen (Figure 2B, Table 1; see supplementary methods section “Multilevel Models for Treatment Effects” for details). The model showed excellent MCMC convergence across all parameters (Supplementary Fig. S2) and suggested that vaccine responsiveness for all regimens but especially DA, i.e. a single early immunisation followed by an adjuvant only boost, was substantially lower in the wild (Figure 2B, Table 1).

However, this model does not explain how habitat and diet are driving poor vaccine responsiveness. Further, it does not account for potential confounding and mediating effects of mouse sex, reproductive status, body mass, body fat, or gastrointestinal parasite burdens, which we hypothesise also contribute to variation in vaccine responsiveness. To address these limitations, we constructed a structural causal model (SCM) of the processes generating variation in vaccine responsiveness ℂ_*VE*_ in this system (See Model Construction and Validation).

### Updating the structural causal model of vaccine responsiveness in light of the data

We tested the validity of the causal hypotheses represented in the DAG 𝔾_*VE*_ using the observed data (Fig 3; see methods sections “Model construction” and “Model Construction and Validation”). For example, 𝔾_*VE*_ (Figure 3A) indicates that parasite infection should be independent of vaccination if adjusted for diet, reproductive status, sex, and habitat (*P* ╨ *V* ∣ *D*, *R*, *S*, *H*); therefore, assuming linearity, we fitted a binomial generalised linear mixed model *P* ∼ *α* + *β*_*V*_*V* + *β*_*D*_*D* + *β*_*R*_*R* + *β*_*S*_*S* + *β*_*H*_*H* + (1|*ID*), and confirmed that the posterior distribution of *β*_*V*_ was not significantly different from zero. This process was repeated for all implied independencies derived from the causal graph 𝔾_*VE*_ (see “Model validation” methods), and 𝔾_*VE*_ was updated until all causal paths were both supported by the observed data and consistent with biological expectations. The final DAG 𝔾_*VE*_ and corresponding structural equations ℂ_*VE*_ are shown in Figure 3.

**Fig. 3:**
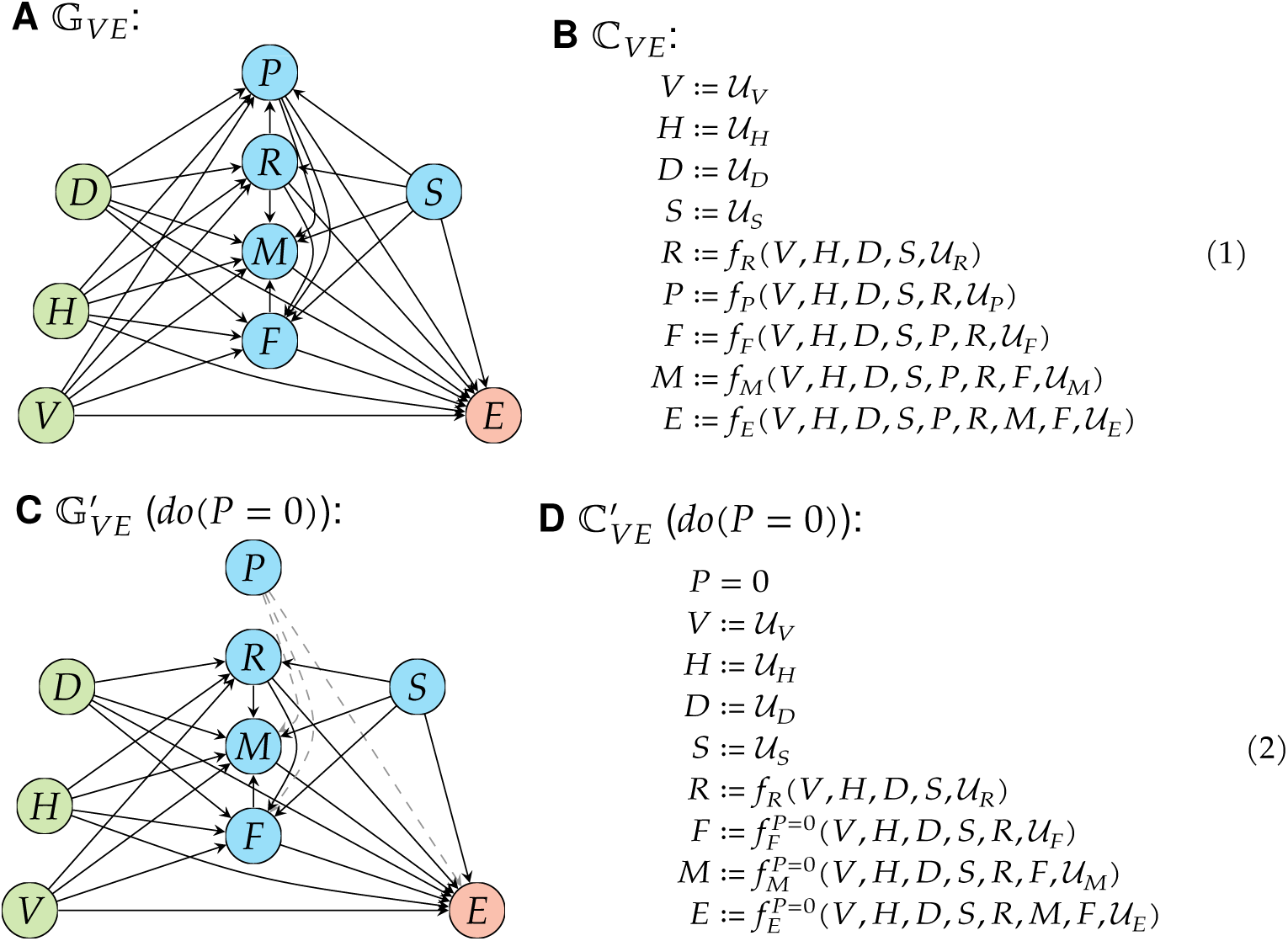
Causal Model of vaccine responsiveness. Structural causal models comprise a causal graph (A, C) and an associated set of structural equations (B, D) for which the functional form is not prescribed (it need not be linear). **A**, Directed acyclic graph 𝔾_*VE*_ representing hypothesised causal effects driving variation in efficacy *E* of vaccine *V* in laboratory or wild wood mice *H*, under supplemented or control diet *D* (green nodes depict experimental treatments). Covariates, mediators, and confounders (blue nodes) included gastrointestinal parasite infection (presence or burden) *P*, reproductive status *R*, body mass *M*, fat scores *F*, and sex *S*. Directed edges depict causal effects hypothesised to operate between nodes. **B**, Structural causal model ℂ_*VE*_ is derived from 𝔾_*VE*_ where each endogenous variable is expressed as a function of its direct causes and an unmeasured error term 𝓊. The functions *f*, *g*, *h*, *i*, and *j* represent the causal mechanisms, with their specific forms left unspecified. **C**. Modified DAG 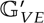 representing the simulated anthelmintic intervention *do*(*P* = 0), where incoming causal paths to *P* are blocked by the intervention. **D**. Structural causal model 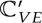 derived from 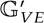 under the intervention, where *P* is set to 0 and its incoming causal paths are removed.

### Causal analysis demonstrates that diet and habitat strongly impact vaccine respons-iveness

Using the resulting validated DAG 𝔾_*VE*_, we fit the set of structural equations ℂ_*VE*_ (Model 1) as Bayesian hierarchical linear models to estimate the total and direct causal effects indicated by the edges in 𝔾_*VE*_ (Fig. 4). All Bayesian models showed excellent convergence with well-behaved MCMC chains and *R̂* < 1.01 for all parameters (See supplementary information “Model Validation”; Supplementary Fig. S3). The posterior distributions of causal parameters (regression coefficients) indicated that vaccination had an average causal effect of 1.50 ± 0.13 (*log*10 + 1)-transformed OD units of DT-specific IgG1 compared to unvaccinated animals. Additionally, the effect of vaccination on antibody production was strongly influenced by both wild habitat and dietary supplementation. However, while body fat *F*, body mass *M*, and reproductive status *R* were also strongly affected by diet and habitat, they did not mediate those effects on vaccine responsiveness.

**Fig. 4:**
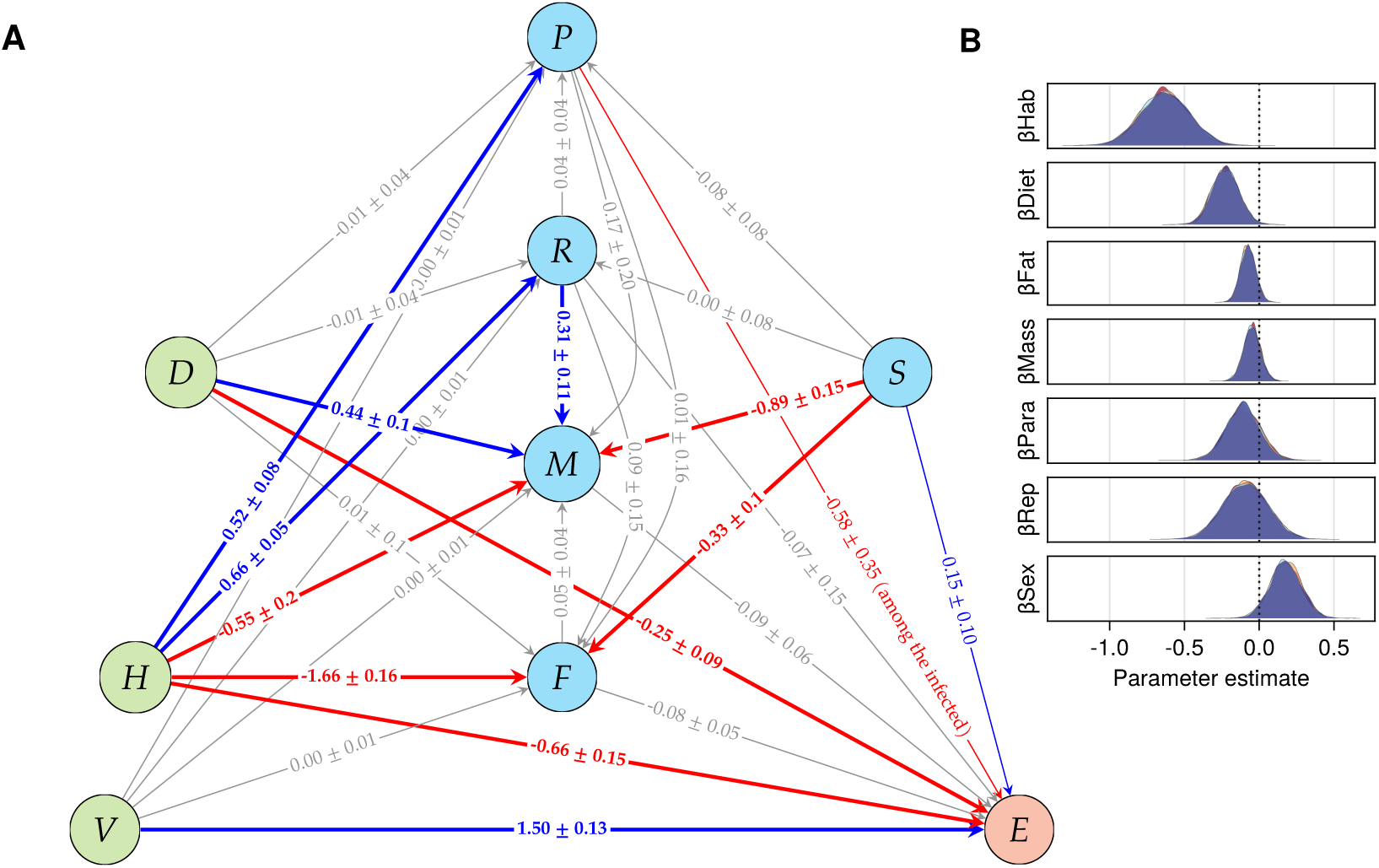
Fully resolved Causal Model of vaccine efficacy. **A**, Directed acyclic graph representing point estimates of positive (blue), negative (red), and null (gray) direct causal effects ± std of Habitat *H*, Diet *D*, Fat scores *F*, body mass *M*, reproductive status *R*, sex *S*, and parasite burden *P* driving vaccine responsiveness *E*. Edges show the direct causal effect of its parent node on its child node conditional on the specific adjustment set for that model. **B**, MCMC sampling of posterior distributions of the coefficients estimating the direct effect of the wild habitat *H* on vaccine responsiveness *E*, with the adjustment set including Diet *D*, Fat scores *F*, body mass *M*, parasite burden *P*, active reproductive status *R*, and female sex *S*; vaccine formulation *V* and individual ID were included as random effects 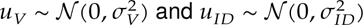, respectively, such that 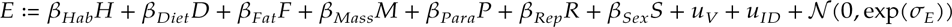 (see Model parameterisation and counterfactual inference methods).

### Impact of wild habitat on wood mice reproduction infection and vaccine responsive-ness

Wood mice in the wild, unlike those in the lab, could reproduce freely and were naturally exposed to the gastrointestinal nematode *Heligmosomoides polygyrus*. As a result, 66% of the wild wood mice were classified as reproductive (R), and 52% were infected with the gastrointestinal nematode *Heligmosomoides polygyrus* (P). Living in the wild also had an overall negative effect on vaccine-specific IgG1 of −0.46 ± 0.24 *logOD*, corresponding to a ∼40% drop in vaccine responsiveness, and a direct negative impact of −0.66 ± 0.15 *logOD* once all other mediating factors were adjusted for (Fig. 4A).

### Diet supplementation reduced vaccine responsiveness

The total causal effect of high quality diet supplementation on vaccine responsiveness was estimated to be −0.25 ± 0.09 *logOD*, adjusted for habitat. This corresponds to an approximate 22% reduction in vaccine responsiveness due to high-quality diet alone. However, the direct effect of diet supplementation on vaccine responsiveness was estimated to be −0.23 ± 0.09 *logOD*, when adjusting for body mass, fat stores, habitat, sex, reproductive status, parasite burden, and vaccine formulation. This indicates that the effect of diet supplementation on vaccine responsiveness was mostly direct (or mediated by unobserved variables along the path *D* → *E*, e.g. immune pathways affecting DTV-specific IgG1 production or the microbiome), but was not mediated by its effects on body mass, fat stores, reproductive status, or parasite burden. Dietary supplementation did increase body mass on average by 2.53 ± 0.3 g, though did not affect body fat scores.

### Sex had direct and indirect effects on vaccine responsiveness

The total causal effect of sex on DTV-specific antibody production *E* indicated greater vaccine responsiveness in females, with approximately an 82% increase in vaccine respons-iveness compared to males (0.26 ± 0.15 *logOD*). However, only a little more than half of the effect of sex (0.15 ± 0.10 *logOD*, ∼ 58% of the total effect) was direct, with body mass (0.08 ± 0.01 *logOD*, ∼ 31% of the total effect) and body fat (0.03 ± 0.005 *logOD*, ∼ 11.5% of the total effect) partly mediating this effect (Fig. 4) as males were both larger and had higher body fat.

### Parasite infection reduced vaccine responsiveness

When considering the entire wood mouse population, including uninfected individuals, *H. polygyrus* burden had no clear effect on vaccine responsiveness (−0.13 ± 0.11 *logOD*). However, because parasite burdens (measured here as the number of adult worms) are overdispersed and zero-inflated (variance/mean ratio = 86.3 for *H. polygyrus*), we also estimated the causal effects of infection burden on DT vaccine responsiveness only among infected mice (Supplementary Fig. S7). Parasite burdens had a strong negative effect on vaccine responsiveness, with a slope of −0.58 ± 0.35 *logOD*, after adjusting for diet, sex, reproductive status, and vaccination. This negative association between parasite infection and vaccine responsiveness was also evident in the raw data analysis (Supplementary Fig. S1B) and indicates that higher parasite burdens are associated with a significant decrease (approx. 74% decrease per 10x parasite count increase) in vaccine responsiveness. Zero-inflated negative binomial models confirmed the robustness of this relationship whilst properly accounting for the ecological reality of uninfected versus infected animals (see Zero-Inflation Modelling for Parasite Data).

### Bayesian prediction of population-wide anthelmintic treatment effects on vaccine responsiveness

To evaluate the potential benefits of anthelmintic treatment for improving Diptheria toxoid vaccine responsiveness, we used Bayesian generative models to simulate the counterfactual scenario where all mice were treated with anthelmintics prior to vaccination. This approach enabled us to predict individual-level vaccine responses under both the observed conditions (with parasites/no drug treatment) and the hypothetical intervention scenario (without parasites due to drug treatment), whilst accounting for all sources of uncertainty in our estimates.

We aimed to (i) quantify the population-wide improvement in vaccine responsiveness after complete parasite elimination, (ii) identify which individuals would benefit most from anthelmintic drug treatment, and (iii) assess the magnitude of the effects of parasite removal in each sex and reproductive cohort. Using posterior predictive sampling from our Bayesian models, we generated counterfactual vaccine responses that maintained each individual’s unique characteristics while removing the effects of parasite infection. Panels C and D of Figure 3 illustrate the structure and equations of the model under the simulated anthelmintic intervention, where all incoming edges to *P* are blocked and *P* is set to zero (Model 2).

Our Bayesian models predicted two distinct but complementary effects of parasite elimination prior to vaccination. At the population level, the overall vaccination coefficient would have improved by 74.3% relative to the population mean among infected individuals (Fig. 5A), representing the enhancement in vaccine responsiveness across the entire population. At the individual level, parasite elimination would have produced a predicted mean increase of 108.1% in vaccine responsiveness per mouse compared to their own individual baseline responses (i.e., with their actual parasite counts). These individual-level benefits varied considerably, ranging from 13.2% to 298.8% improvement (Cohen’s d mean = 0.20 ± 0.03).

**Fig. 5:**
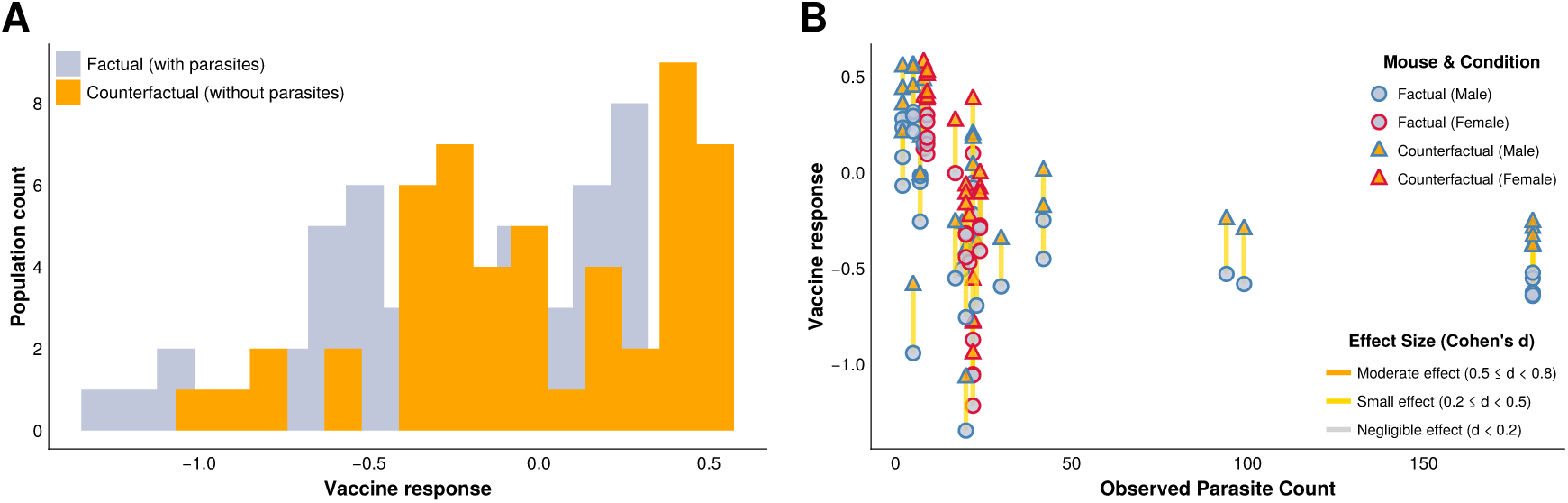
Effect of simulated population-wide anthelmintic drug treatment on vaccine responsiveness. **A**, Population-wide distributions of vaccine responsiveness under observed vs. parasite-free conditions. Density plots comparing vaccine responsiveness in the observed data (with parasites; light grey) versus the counterfactual data (parasite elimination scenario; orange). The rightward shift of the distribution predicts a population-wide improvement in vaccine responsiveness when parasite burdens are eliminated by drug treatment. **B**, Individual-level vaccine response changes following parasite elimination. Each wood mouse’s predicted vaccine responsiveness in factual (circles) versus counterfactual (triangles) scenarios in males (blue outlines) and females (red outlines), plotted against initial parasite burden. Connected lines show individual-level changes, with effect size (Cohen’s d) indicated by the colour of the line.

Further, the magnitude of the predicted improvement in vaccine responsiveness was not strictly proportional to the observed parasite burdens (Fig. 5B), with sex and reproductive status contrubuting to the effect of parasite elimination on vaccine responsiveness (Table 2).

**Table 2:**
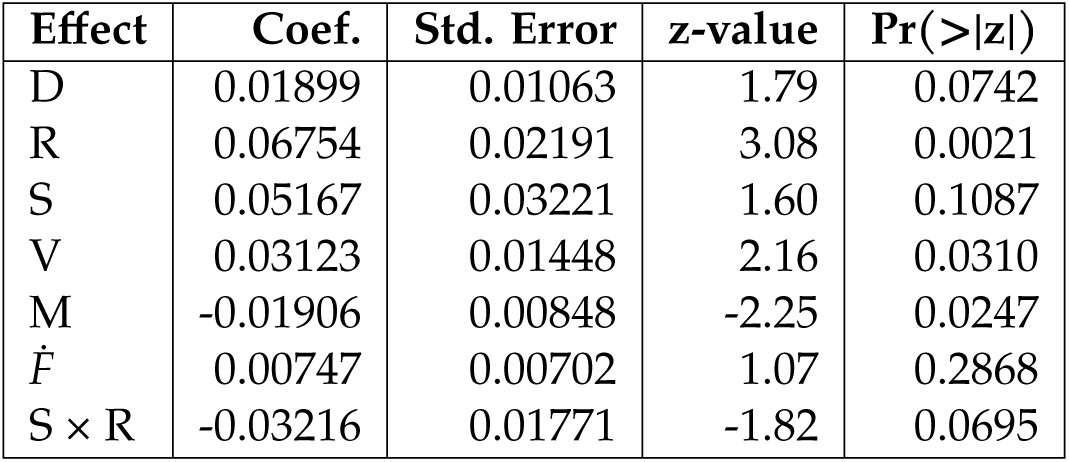
Summary of the linear mixed model estimating the individual-level effect sizes (Cohen’s d) of anthelmintic treatment on vaccine responsiveness. The model includes fixed effects for Diet (*D*), Reproductive status (*R*), Sex (*S*), Vaccination status (*V*), Body mass (*M*), Fat scores (*FḞ*), and the interaction between Sex and Reproductive status (*S* × *R*). Coefficients (Coef.), standard errors, z-values, and p-values (Pr(> |*z*|)) are reported for each effect. This table quantifies the predictors of variation in vaccine response improvements following parasite elimination, highlighting significant contributions of reproductive status, vaccination, body mass, and a marginal interaction between sex and reproductive status.

## DISCUSSION

Using a novel lab-to-wild rodent system in combination with a robust structural causal modelling approach, we provide a test of how key intrinsic and environmental factors impact vaccine responsiveness to a standard Diphtheria toxoid vaccine. We observed substantial variation in vaccine responsiveness between individuals, as measured by vaccine-specific IgG1 antibodies. In addition, wood mice living in natural populations in the wild, had on average 46.9 ± 0.1% lower vaccine-specific antibodies than individuals living in the controlled, more sterile laboratory setting. Countrary to predictions, access to high-quality diet resulted in lower vaccine responsiveness in mice living in both habitats. However, among individuals currently infected with gastrointestinal nematode parasites, there was a negative effect of parasite burdens on vaccine responsiveness. Our results emphasise the importance of accounting for the interplay between environmental and individual factors when analysing and predicting responses to immunisation, consistent with reports across multiple habitats and species, including humans [21, 45, 46, 47].

Indeed, there was marked hyporesponsiveness in the wild wood mouse populations relative to their laboratory conspecifics immunised with the same Diphtheria Toxoid vaccine formulations. Mice living in their natural woodland habitat had significantly lower vaccine responsiveness, and were only partially rescued by a second booster dose of the vaccine. The timing of our measurements (7-35 days post-immunisation) captured both primary and secondary immune responses, revealing that wild mice required booster doses to achieve responses similar to laboratory wood mice after a single dose. The median peak responses occurred 22.0 days post-vaccination (IQR: 14.0-35.0 days) across both habitats; however, the wild wood mice had consistently lower vaccine responsiveness (Supplementary Fig. S6; see Temporal Dynamics Analysis). This suggests that the natural environment may reduce the magnitude of vaccine responses without substantially altering the timing of the immune response, potentially affecting both efficacy and the duration of protection. This reduction in vaccine responsiveness is consistent with widespread reports of vaccine failure during clinical trials, notably in phase III where vaccines are tested for efficacy in naturally exposed individuals [28], or for already-established vaccines when used in rural populations [19, 48].

While we used alum as an adjuvant, the observed differences between habitats raise questions about whether alternative adjuvants might perform better in wild populations. For example, adjuvants that stimulate stronger Type 1 immune responses might be more effective in environments where Type 2 responses are suppressed, as in populations with chronic helminth infections. The effect of adjuvants on vaccine efficacy warrants further investigation, particularly given the growing availability of novel adjuvants that could potentially overcome some of the environmental constraints we observed.

In our study, part of the loss of vaccine responsiveness in the wild habitat was mediated by infection with the gastrointestinal nematode *H. polygyrus*. High worm burdens reducing vaccine responsiveness is consistent with reports that parasitic helminths negatively affect vaccination [10, 49, 50, 51, 52]. These reports are not unanimous, however, with many reporting no [53] or even a positive association between vaccine responsiveness and parasitic infection [11, 13]. While parasite infection can have varied effects on vaccine responsiveness, our study suggests that conflicting inferences might be due to how the effects of parasite infection are estimated. Specifically, we only detected a negative effect of parasite burden among the infected animals, and no effect when uninfected individuals were included in the analysis. Thus, we might predict that vaccination campaigns that include anthelmintic treatment as part of their regime, could lead to detectable population-wide improvements in vaccine efficacy for populations where nematode infection prevalence and burdens are high. The observed variation in individual responses, along with the high prevalence of parasite infections in rural populations, suggests that achieving herd immunity may require higher vaccination coverage than typically assumed. This finding could significantly impact vaccination strategies in natural populations, especially in areas where parasite infections are endemic. Our models indicate that reproductive females are likely to benefit the most from anthelmintic treatment, particularly in improving vaccine responsiveness.

While we were unable to investigate the mechanisms by which *H. polygyrus* affected vaccine responsiveness, two hypotheses have been proposed. The first suggests counter-regulation between different arms of the immune system, e.g., Th2 responses might suppress pathways that drive vaccine-specific antibody production[54, 55], while the second hypothesis is focused on immune suppression by the parasites [27, 56]. Here, the immune system of wood mice naturally infected by *H. polygyrus*, which, while distinct from the laboratory model [57], might similarly be actively suppressed by the parasite. Alternatively, animals inherently more susceptible to helminth infection might have a lower propensity to generate strong antibody responses, which we detect as poor vaccine responsiveness. Resolving these alternatives would require testing the prediction of both vaccine responsiveness and nematode burden from immune profiling before both infection and vaccination (see for example [58, 59]).

While our study focussed on nematodes, other infections are likely to have impacted vaccine responsiveness in the wild, including viruses, bacteria, and other helminths. In particular, reports that variability in responses to SARS-CoV2 in humans could be explained by cytomegalovirus infections [14], and given the 10–70% prevalence of wood mouse herpesvirus infections in our study populations [60], it is likely that this virus (or other parasites/pathogens) could have also impacted vaccine responsiveness in our study.

High-quality dietary supplementation had unexpected negative effects on vaccine responsiveness, consistently reducing vaccine-specific antibody responses in both wild and laboratory populations. Several explanations may account for this counter-intuitive result. First, high-quality nutrition may shift immune investment towards Th2-dominated responses and tissue repair functions, at the expense of inflammatory and Type 1 pathways that support vaccine-specific antibody production. This could reflect evolved trade-offs in immune resource allocation under different nutritional states. Our experiment (July-September) coincided with peak natural food availability, potentially creating a context where wood mice with access to high-quality supplemental diet experienced nutritional excess rather than relief from scarcity. The effects of dietary supplementation may also be context-dependent, with benefits primarily occurring under natural food limitation. Dietary changes may also alter gut microbiome composition in ways that affect systemic immune responses. High-quality laboratory chow may promote microbiota that are less conducive to vaccine responsiveness compared to diverse natural diets. Additionally, the alum adjuvant used may interact differently with metabolic pathways activated by high-quality nutrition, potentially dampening rather than enhancing the intended immune response. Finally, the observed effect may reflect unmeasured confounding or specific characteristics of our study population and timing, rather than a universal biological phenomenon.

Our previous research in both the wild and laboratory setting found that wood mice with access to this high-quality diet had ∼50% reductions in *H. polygyrus* egg and worm burdens and that anthelmintic treatment was more effective [40]. In this study, however, we didn’t detect an effect of food supplementation on parasite burdens. This discrepancy could result from: (i) seasonal variation in natural food availability eliminating the relative benefit of supplementation; (ii) different parasite exposure pressures between study periods; or (iii) non-linear dose-response relationships where benefits occur only under specific nutritional stress conditions.

Next, while our structural causal model provides a principled framework for causal inference, several limitations affect our confidence in causal claims regarding diet effects: (i) the assumption of no unmeasured confounding effects may be violated by unobserved variables such as microbiome composition, stress responses, or behavioural changes; (ii) the consistency assumption may not hold if “dietary supplementation” represents a complex intervention affecting multiple biological pathways simultaneously; and (iii) temporal confounding due to seasonal effects cannot be completely ruled out. These findings highlight the critical need for mechanistic studies to elucidate how nutrition affects vaccine responses across different contexts. Future investigations should include: immune pathway profiling (Th1/Th2 balance, cytokine responses), microbiome analysis, dose-response studies, and experiments across different seasons and nutritional baselines. Without such mechanistic validation, our dietary findings should be interpreted cautiously.

Females overall had higher vaccine responsiveness than males, partly as a direct effect of sex differences and partly mediated by body mass and fat content. Sexual dimorphism in disease susceptibility and immunity has been widely reported across multiple host and pathogen species, with females typically more resistant to infection including by helminths [61], generating stronger responses to vaccination, but more likely to develop autoimmunity [8, 62, 63, 64, 65]. Large-scale studies have identified sexual dimorphism in specific immune pathways, with females tending to present higher activation of innate pathways [66], though this might be especially true in younger individuals [67]. In our study, mice tended to be in the early adult stage, possibly skewing younger in the wild cohort where lifespans are likely to be much shorter, though this age effect was likely overshadowed by other negative effects on vaccine responsiveness. After adjusting for habitat, while females had lower body mass and fat stores, these differences had no effect on DTV-specific antibody responses. In fact, we only detected a direct effect of sex, and no indirect effects.

Finally, Bayesian generative models predicted vaccine responsiveness under hypothetical population-wide anthelmintic treatment, addressing two complementary questions. We estimated that eliminating parasites would have improved the overall vaccination coefficient by 74.3% among infected individuals, but only 4.3% when averaged across the entire population (including uninfected individuals). This population-level metric reflects the policy-relevant question: “How much better would vaccines work overall if we implemented mass anthelmintic treatment?” The modest population-level improvement occurs because only 52% of the wild population was infected, the effect is diluted when averaged across infected and uninfected individuals, and population-level estimates reflect marginal effects averaged across all individuals.

However, several assumptions underlying our counterfactual analysis warrant caution: (i) No interference assumption: We assumed treating one individual does not affect others, but anthelmintic treatment could reduce transmission and benefit untreated individuals through herd effects; (ii) Intervention feasibility: Our model assumes perfect parasite elimination, whereas real anthelmintic treatments have variable efficacy and may require repeated dosing; (iii) Temporal considerations: We modelled instantaneous para-site removal, but actual treatment effects unfold over time and may interact with immune recovery dynamics.

Despite these limitations, our results suggest that anthelmintic treatment could provide meaningful benefits for increasing vaccine responsiveness in helminth-endemic populations, consistent with human studies [51, 10, 50, 52]. The substantial individual-level heterogeneity (20-fold variation in benefit) indicates that targeted approaches identifying high-burden individuals could be more cost-effective than mass treatment programmes, though this would require practical diagnostic tools for parasite burden assessment.

Importantly, our study was limited to variables that could be measured in a wild, non-model, species. Further studies are needed to disentangle how different arms of the immune system mediate the effects of vaccination on antibody production, and how the environmental factors may modulate those different pathways. We also focussed on a single antibody isotype, IgG1. While this is appropriate for assessing alum-adjuvanted vaccines, further studies should investigate protective immunity to specific pathogens after vaccination. Indeed, while our conclusions are consistent with reports that investigate protective immunity [19], it is likely that inferences would differ depending on the pathogens targeted and the vaccination formulations employed. The use of optical density measurements for quantifying vaccine-specific IgG1, while standard in vaccine studies, might not capture all aspects of antibody functionality. Future studies could incorporate additional measures of antibody quality, such as affinity maturation and neutralisation capacity, to provide a more comprehensive understanding of vaccine-induced protection. Beyond the immune system. It might also be useful to explore how the gut microbiota might mediate the dietary effects, and how different nutrients, quality, and abundance, whether supplied or available in the environment, might drive the interactions that impact vaccine responsiveness.

Lastly, taken together, this study provides a useful methodological approach to es-timate causal relationships between environmental and individual factors and vaccine responsiveness. By employing explicit causal inference rather than conventional statistical approaches that can obscure true causal mechanisms [39], this approach could be used to predict vaccine responsiveness in target populations, and to identify which popula-tions or subsets are likely to respond poorly to a specific vaccine, and/or benefit from targeted interventions such as diet changes, antiparasitic treatments, or different vaccine adjuvants/regimes. This could be particularly useful for vaccines that are known to be sensitive to environmental factors, such as anthelmintic vaccines, and for which the causes of vaccine hyporesponsiveness remain difficult to identify, quantify, and therefore, address. Our study also indicates that laboratory studies may not always be generalisable to wild populations, and are likely to systematically overestimate vaccine efficacy. This suggests that the development of vaccines should be more closely aligned with the conditions in which they are intended to be used, and that the effects of environmental factors on vaccine responsiveness should be more thoroughly investigated.

## MATERIALS, METHODS, AND MODELS

All animal work was conducted in compliance with the UK Home Office Animals (Sci-entific Procedures) Act, 1986. All laboratory and field experiments were approved by the University of Edinburgh Ethical Review Committee and carried out under the UK Home Office Project Licence 70/8543. The dosage of diphtheria vaccine was tested in laboratory-bred wood mice before the experiments for safety. Animal sacrifice was per-formed using appropriate Schedule 1 Methods. Fieldwork was carried out with permission of the Forestry Commission Scotland under the permit SUR09.

### Vaccine formulation and administration

The vaccine used in this study was a commercially available diphtheria inactivated toxoid (DT, Alpha Diagnostic International). The vaccine, DTV henceforth, was prepared in our laboratory one day before administration, using 2 Lf of vaccine-grade DT protein adsorbed to the adjuvant (9% alum). The vaccine was administered subcutaneously in a volume of 100μl. The control group was injected with 100μl of adjuvant (9% alum) suspended in 1X PBS.

### Laboratory wood mouse experiments

#### Wood mouse colony and housing

The laboratory experiments were carried out on a colony of wild-derived but now captive outbred wood mouse colony bred and maintained at the University of Edinburgh [40]. The laboratory wood mouse colony has been maintained as an outbred colony for over 10 generations. Mice were housed in conventional laboratory conditions with controlled temperature (20-22 °C), humidity (45-65%), and 12:12 light-dark cycle. Standard husbandry followed UK Home Office guidelines with enrichment including nesting material and shelter. All laboratory mice used in experiments were 8-16 weeks old and sexually mature but reproductively naive.

#### Diets

Two commercial laboratory diets were used in this study. TransBreed^TM^ (SDS Diets Ltd., UK) is a high-quality breeding diet containing 20.1% crude protein, 10.1% crude fat, 3.5% crude fibre, and 4.9% crude ash, with enhanced nutritional content optimised for reproductive performance and immune function. We have previously shown that wood mice supplemented with TransBreed^TM^ are more resistant to the gastrointestinal nematode *Heligmosomoides polygyrus*, cleared worms more effectively after anthelmintic treatment, and produced stronger general (total IgA) and parasite-specific (IgG) immune responses in both wild and laboratory conditions [40]. The other half of the mice in this experiment were fed RM1. RM1 (Rat and Mouse Maintenance 1, SDS Diets Ltd., UK) is a standard maintenance diet containing 14.4% crude protein, 2.7% crude oil, 4.7% crude fibre, and 6.0% crude ash. TransBreed^TM^ provides substantially higher energy density (20.1% vs 14.4% protein; 10.1% vs 2.7% fat) and enhanced micronutrient content compared to RM1, including elevated levels of vitamins, minerals, and essential fatty acids that support improved immune function and reproductive performance. Both diets were used in the laboratory colony, but only TransBreed^TM^ was used for supplementation in the wild experiment.

#### Vaccination regime

The laboratory experiment was conducted in two replicate blocks, with 36 animals in each block (18 males, 18 females). At 12 days before the start of the experiment (d-12), all mice were shifted to new diet regimes and given time to acclimatise; half of the animals in each block (9 males and 9 females) were given TransBreed^TM^. Within each diet regime, both male and female mice were randomly assigned to one of three experimental groups (Fig. 1). On d0, animals belonging to Groups ‘DD’ & ‘DA’ (n = 24) were subcutaneously injected with 100μl of DTV, and Group ‘AD’ (n = 12) were injected with the adjuvant control. Small blood samples were taken via tail snip on d14 and d17 to measure the primary antibody response. Twenty-one days (d21) after the first injections, the animals of Group ‘DD’ (n = 12) were injected again with 100μl of DTV (‘DD’: diphtheria vaccine followed by diphtheria booster), while those assigned to Group ‘DA’ (n= 12) were injected with the control dose as described above (‘DA’: DTV followed by adjuvant only). Animals assigned to Group ‘AD’ (n = 12) that had previously been injected only with the control adjuvant only dose now received an injection of 100μl DTV (‘AD’: adjuvant followed by DTV). Another blood sample (20-50μl) was taken a day later (d22) by cheek venipuncture. All animals were sacrificed on d35 (i.e., 14 days after the second injection) and a blood sample was taken for measuring the antibody response. The same procedures and timeline were carried out for the second experimental block of animals (n = 36). Mice were co-housed throughout the experiment, in same-sex groups of three, with equal representation of the three experimental groups in each cage to prevent confounding cage effects.

#### Wild wood mice experiment

We conducted a 10-week field experiment in a natural population of wood mice with a design similar to that of the laboratory experiment described above. The experiment was conducted in a woodland in Falkirk, Scotland, UK (Callendar Wood, 55.990470, −3.766636), where we established four trapping grids (60m × 40m per grid) with 10m spacing between each of the 35 trapping stations per grid. At each station, we set a pair of Sherman live traps (H.B. Sherman 2 × 2.5 × 6.5-inch folding trap, Tallahassee, FL, USA). We randomly selected two of the four grids for dietary supplementation, initiating the treatment 12 days before live-trapping began. Specifically, 6kg of TransBreed^TM^ pellets were evenly scattered across each treated grid twice weekly (approximately 170g per trapping station), with supplementation continuing throughout the experiment. The remaining two grids served as controls and received no additional food. All wood mice had access to their natural diet, so the supplemental pellets on the treated grids provided ad libitum access to high-quality nutrition while allowing continued foraging for natural food sources.

From July-September 2018, wood mice were trapped 2–3 nights per week using live traps baited with grains, carrots, and bedding. Additionally, traps on the supplemented grids were also baited with 1-2 TransBreed^TM^ pellets. For identification, all newly captured mice weighing >13g were subcutaneously tagged with a unique 9-digit microchip passive induced transponder (PIT tag; FriendChip AVID2028, Norco, CA, USA). Upon first capture, mice were randomly assigned to one of the three vaccine treatment groups ‘DD’, ‘DA’, and ‘AD’, described above and in Fig. 1; mice assigned to these groups received the same primary and booster doses of DTV or alum control as described above. Throughout the experiment, small volume blood samples were taken weekly via tail snip from each tagged animal to measure their antibody response to the diphtheria immunisation, on days as close to possible as their laboratory counterparts. Animals recaptured 14 or more days after their second injection were sacrificed and a terminal bleed was collected to measure their antibody response and adult *Heligmosomoides polygyrus* worm burdens. *H. polygyrus* is a natural gastrointestinal nematode found at high prevalence (20-100 %) in wild populations of *A. sylvaticus* [40, 68, 69, 70], as well as being a well-studied model system of human gastrointestinal nematodes where it has been found to be highly immunomodulatory [56, 71].

Additionally, at all captures, we took the following demographic metrics for each individual: sex, body condition, reproductive condition, weight (g), and length (mm). Sex and reproductive condition of each mouse was assigned by examining the genitals, with males having larger urogenital gap compared to females. Males were classified as reproductively active if their testes had visibly descended to scrotal sacs, and females were classified as reproductively active if they had a perforated vagina or were visibly pregnant or lactating. Body condition was measured by assigning dorsal and pelvic fat scores on a qualitative scale from 1-5, where 1 represented a mouse with very low condition/fat reserves, while a 5 represented a mouse with ample fat reserves. This was achieved by palpating the back and pubic bones [72]. Both scores were added to provide a single metric of body condition for analysis.

#### Vaccine responsiveness: DT-specific IgG1

Vaccine responsiveness was measured as the concentration of anti-diphtheria IgG1 antibodies. This is the gold standard method used in vaccinology to measure the antibody-specific response to immunisation and is used specifically for the diphtheria toxoid vaccine used here [73]. Blood samples from both the laboratory and the wild mice were centrifuged @24,000 rpm for 10 min within 4–6 hours of collection. The sera were separated from the pellets and stored at −80 °C until further analysis. 96-well plates (NuncTM MicroWellTM) were coated overnight at 4 °C with diphtheria toxoid (2μg/mL) diluted in carbonate buffer (50 μl per well). After washing the plates with TBS (10 X) and Tween80 thrice, 100 μl of TBS(1X)-4% BSA was added per well and incubated at 37 °C for 2 hours to block non-specific binding sites. Plates were then tapped dry and 50 μl of serum samples serially diluted (starting dilution 1:100) in TBS (1X)-4% BSA buffer were added per well and left at 4 °C for binding overnight. The next day, 50 μl HRP-conjugated anti-mouse IgG1 detection antibody (Southern BioTech) per well were added after washing the plates 4 times with TBS (10X) and Tween80 and tapping them dry. After incubation at 37 °C for 1 hour, the plates were washed again, four times with TBS (10X) and Tween80 and then twice with dH20. Then, 50 μl of TMB substrate solution were added per well and the enzymatic reaction was left to develop in the dark for 7 min. The reaction was stopped after 7 min using 50 μl sulphuric acid (0.18 M) per well. Absorbance at 450 nm was recorded using an ELISA plate reader (Multiscan, Ascent Labsystems) immediately thereafter. For each sample, blank-centred optical densities (OD) of three consecutive wells (of dilutions 1:3200, 1:6400, 1:12800) were averaged to obtain diphtheria-specific IgG antibody measurements for each individual.

#### Data processing and analyses

ELISA optical densities of diphtheria (DT)-specific antibodies (*E*) were log_10_(1 + *x*)-transformed to improve normality, while body weight (*M*) and fat scores (*F*) were standardised (Z-score transformed) to facilitate prior specification and improve computational stability. All other variables were treated as binary. All data processing was performed in Julia v1.11 [74] using packages CSV.jl [75] and DataFrames.jl [76]. Large Language Models were used for code refactoring and optimisation. Delailed quantitative methods are available in the supplementary methods section “Bayesian Statistical Implementation”. All code is available at https://github.com/SimonAB/Apodemus_vaccines

#### Generalised linear mixed models of the effects of experimental interventions

We used generalised linear mixed models (GLMMs) to estimate the main effects of habitat, diet, and vaccine formulation on vaccine-specific antibody concentrations. Mouse ID was modelled as a random effect to account for repeated measurements. Interactions between vaccine regime and habitat, and between diet and habitat, were included to test whether wild mice responded differently to treatments than laboratory mice. Likelihood ratio tests (LRT) evaluated the contribution of interaction terms. The package MixedModels.jl [77] was used for all GLMMs and LRTs.

We then implemented Bayesian hierarchical models to minimise the effects of data imbalance and provide robust uncertainty quantification. Weakly informative Gaussian priors were specified for intercepts and regression coefficients. Residual standard devi-ations were given Exponential(1) priors, and random effect standard deviations used Exponential(1) priors with non-centred parameterisation to improve sampling efficiency. Prior predictive checks are performed to ensure biological plausibility (Supplementary Fig. S4). Posterior estimates were sampled using Hamiltonian Monte Carlo with the No U-Turn Sampler [78], with 4 chains of 3,000 iterations each after 1,000 warmup iterations. Convergence was assessed using *R̂* < 1.01 for all parameters and effective sample sizes > 400. Turing.jl [79] was used for Bayesian modelling; Makie.jl v0.17 [80] was used for plotting.

#### Structural Causal Models

Structural causal modelling (SCM) [42] provides a framework for identifying and estimating causal effects from observational data by explicitly representing causal assumptions through directed acyclic graphs (DAGs). Prior predictive checks validated that our weakly informative priors produced biologically plausible vaccine responses whilst appropriately covering the observed data space (Supplementary Fig. S4). We used SCMs to quantify the causal pathways through which habitat, diet, and parasite infection influence vaccine responsiveness, distinguishing between direct effects (e.g., parasites directly suppressing immune responses) and indirect effects mediated through body condition or reproductive status.

Our primary estimand was the *average direct causal effect of wild habitat on vaccine respons-iveness* conditional on diet, sex, reproductive status, body mass, fat scores, and parasite burden: 𝔼[*E*^*H*=1^ − *E*^*H*=0^ ∣ *D*, *S*, *R*, *M*, *F*, *P*], where *H* = 1 denotes wild habitat and *H* = 0 denotes laboratory habitat. Secondary estimands included the direct causal effects of diet supplementation, parasite burden, and sex on vaccine responsiveness, and the *average treatment effect of hypothetical anthelmintic intervention*: 𝔼[*E*^*P*=0^ − *E*^*P*=*observed*^], representing the population-level improvement in vaccine responsiveness under complete parasite elimination.

Our approach relied on several key identifying assumptions:

1. **No unmeasured confounding:** We assumed no unobserved variables jointly affect both exposures and outcomes after conditioning on measured covariates.
2. **Stable Unit Treatment Value Assumption (SUTVA):** We assumed each individual’s response depends only on their own treatment assignments, not others’.
3. **Consistency:** We assumed that “dietary supplementation” and “wild habitat” represent well-defined, consistent interventions.
4. **Linearity:** We assumed relationships between variables are approximately linear on the transformed scales used.
5. **Positivity:** We assumed sufficient overlap in covariate distributions across treatment groups to enable causal inference. While randomisation ensures this for assigned treatments, natural variation in parasite burden, body condition, and reproductive status may create sparse regions of covariate space where extrapolation is required.

To assess robustness to assumption violations, we implemented comprehensive sensitivity analyses including: (i) E-value calculations [81] to quantify the minimum strength of unmeasured confounding needed to explain away observed effects (Supplementary Fig. S8; see E-value Sensitivity Analysis for Unmeasured Confounding); (ii) prior sensitivity analysis to evaluate robustness to model specification across different prior scales (see Prior Sensitivity Analysis); (iii) zero-inflation modelling for realistic parasite count analysis (see Zero-Inflation Modelling for Parasite Data); (iv) temporal dynamics analysis to characterise vaccine response kinetics (see Temporal Dynamics Analysis); and (v) causal assumption testing beyond standard conditional independence (see Causal Assumption Testing).

#### Model construction

We constructed the causal model through an iterative process of hypothesis formulation, graphical representation, and empirical validation (see section “Model validation” and causal flow diagram, Fig. S5). The DAG was built using domain knowledge and experimental design constraints:

Randomised treatments (vaccine *V*, diet *D*, habitat *H*) have no parent nodes. Sex *S* is exogenous due to random allocation across treatments. We hypothesised that habitat affects all measured variables except vaccine and sex; diet affects reproductive status *R*, body mass *M*, body fat *F*, parasite burden *P*, and vaccine response *E*; and fat reserves *F* causally precede body mass *M*. Vaccine responsiveness *E* was modelled as affected by all other variables. Each variable includes an independent error term capturing unmeasured sources of variation.

Model construction was performed in Julia v1.11 [74] using Dagitty.jl for causal identification and Turing.jl [79] for Bayesian estimation. The DAG 𝔾_*VE*_ corresponding to ℂ_*VE*_ was visualised using Tikz [82].

#### Model validation

We validated the structural causal model ℂ_*VE*_ (Model 11) by testing the conditional independence relationships it implies [42, 83]. The DAG entails 13 conditional independence statements: (*D* ╨ *S*), (*D* ╨ *V*), (*H* ╨ *S*), (*H* ╨ *V*), (*S* ╨ *V*), (*F* ╨ *V* ∣ *D*, *H*), (*M* ╨ *V* ∣ *D*, *H*), (*R* ╨ *V* ∣ *D*, *H*), (*P* ╨ *V* ∣ *D*, *H*), and (*P* ╨ *V* ∣ *D*, *R*, *S*, *H*). We tested each independence using generalised linear mixed models with appropriate link functions, including mouse ID and immunisation protocol as random effects. Independence statements were considered supported when *P* > 0.05, providing conservative validation of the causal structure. MixedModels.jl [77] was used for conditional independence testing. All 13 independence tests were satisfied, validating the proposed causal structure.

#### Model parameterisation and counterfactual inference

Bayesian models were fitted using the same prior specifications as the hierarchical models above, with the structural equations encoded as a system of regression models. All modelling was performed in Julia v1.11 [74] using Turing.jl [79].

##### Statistical identification

The DAG 𝔾_*VE*_ was encoded as a Bayesian network representing each variable as a conditional distribution given its parents. Posterior distributions were estimated using Hamiltonian Monte Carlo with the No U-Turn Sampler [78], with 4 chains of 3,000 iterations each after 1,000 warmup iterations. Convergence diagnostics included *R̂* < 1.01, effective sample sizes > 400, and visual inspection of trace plots.

##### Missing data imputation

Twenty-two mice had missing fat scores (67 missing values total). We used Bayesian imputation within the structural model, where missing values were modelled as draws from their conditional distribution given observed par-ent variables: *Ḟ*_*missing*_ ∼ Normal(*v*_*F*_, *σ*_*F*_), where *v*_*F*_ and *σ*_*F*_ were estimated from the data (see supplementary methods section “Bayesian Generative Models for Counterfactual Inference” for details).

##### Counterfactual estimation of anthelmintic treatment

To evaluate potential benefits of anthelmintic treatment, we implemented the *do*-calculus [42] to simulate the intervention *do*(*P* = 0) (complete parasite elimination). Prior predictive checks confirmed that both factual and counterfactual models were well-calibrated with appropriate prior coverage (Supplementary Fig. S4). This involved fitting two Bayesian models: the factual model including all causal pathways, and the counterfactual model with the parasite effect on vaccine response set to zero while maintaining all other pathways (Figure 3C-D, Model 20). For each mouse, we generated posterior predictive samples under both scenarios, calculating individual treatment effects as the difference between counterfactual and factual predictions. Effect sizes were calculated using Cohen’s *d* with pooled standard deviations, categorised as negligible (|*d*| < 0.2), small (0.2 ≤ |*d*| < 0.5), moderate (0.5 ≤ |*d*| < 0.8), or large (|*d*| ≥ 0.8). This approach quantified both population-level improvements and individual heterogeneity in treatment benefits while propagating all sources of uncertainty through the causal model.

We calculated intervention effects using two complementary approaches: (1) **Population-level vaccination effect improvement**, which compares vaccination coefficients (*β*_*V*_) between factual and counterfactual scenarios using mixed-effects models, quantifying how much better vaccines work overall when parasites are eliminated; and (2) **Individual-level response improvement**, which calculates the percentage change in each mouse’s predicted vaccine response between factual (with parasites) and counterfactual (without parasites) scenarios. The population-level metric reflects policy-relevant vaccine efficacy changes, whilst individual-level metrics capture the magnitude of benefit each animal would experience from parasite elimination. (See additional details in supplementary material “Effect Size Calculation and Clinical Significance”.)

While it would have been desirable to test interactions between sex and reproductive status, sample size limitations allowed only for testing the main effects of sex and reproductive status on vaccine responsiveness under the counterfactual scenario (see supplementary methods section “Bayesian Generative Models for Counterfactual Inference” for full details).

## ACKNOWLEDGEMENTS

This work was supported by a PhD studentship from the Darwin Trust of Edinburgh for SV and ARS, as well as Wellcome Trust Institutional Strategic Support Fund (ISSF) grants to ABP (ISSF 2014; J22737) and SAB (097821/Z/11/Z). Additionally, SAB received a targeted School of Biodiversity, One Health, and Veterinary Medicine Research Fellowship, and ABP was awarded a University of Edinburgh Chancellor’s Fellowship. NERC grant NE/X01424X/1 also supported ABP and SAB. We are grateful to Rivka Lim for her helpful discussions and feedback on the manuscript and modelling.

## AUTHOR CONTRIBUTIONS

Conceived and designed the experiments: ABP, SAB Performed the experiments: SV, JH, ES, AS, ABP, SAB Analysed the data: SV, ES, SAB Contributed reagents/materials/analysis tools: ABP, SAB All the authors reviewed the manuscript.

## SUPPLEMENTARY METHODS

### Structural Causal Modelling Framework

#### Structural Causal Model Equations

The structural causal model ℂ_*VE*_ for vaccine efficacy consists of the following system of equations:

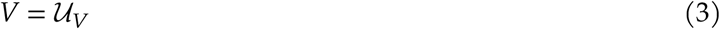

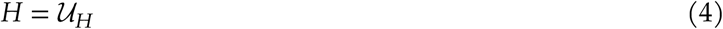

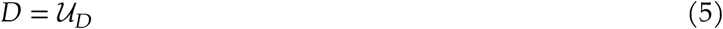

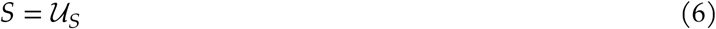

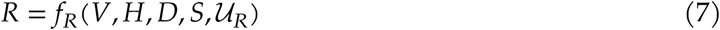

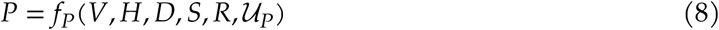

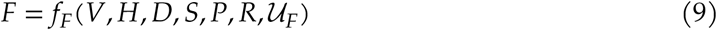

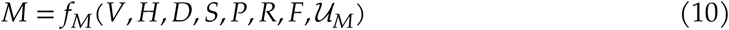

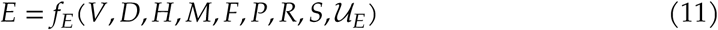

where 𝓊_*j*_ represents independent error terms for each variable *j*. The counterfactual model under the intervention *do*(*P* = 0) sets *P* = 0 and removes its causal effects whilst maintaining all other causal pathways:

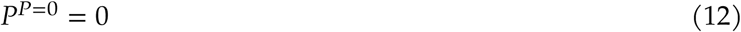

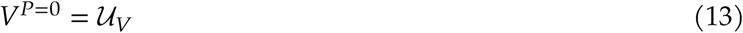

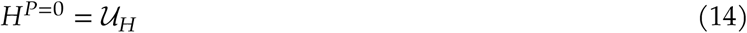

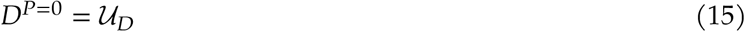

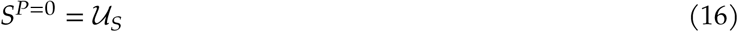

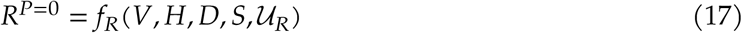

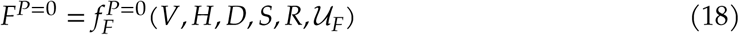

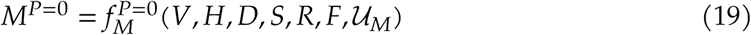

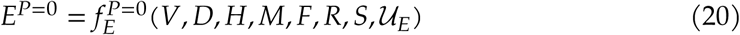

where the functions 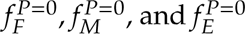 represent the modified causal mechanisms with the direct effects of *P* removed.

##### Model Construction and Validation

Structural causal model construction followed established protocols [42]. Initial DAG construction was based on biological knowledge and experimental design, with treatments (vaccine, diet, habitat) having no parents due to randomisation. The causal structure was refined through iterative testing of conditional independence statements implied by the graph.

The validity of the structural causal model was assessed by testing 13 conditional independence statements implied by the DAG using generalised linear mixed models. Each independence statement was tested at the *α* = 0.05 significance level to provide conservative validation:

1. *D* ╨ *S* (Diet independent of Sex)
2. *D* ╨ *V* (Diet independent of Vaccination)
3. *H* ╨ *S* (Habitat independent of Sex)
4. *H* ╨ *V* (Habitat independent of Vaccination)
5. *S* ╨ *V* (Sex independent of Vaccination)
6. *F* ╨ *V* ∣ *D*, *H* (Fat scores independent of Vaccination given Diet and Habitat)
7. *M* ╨ *V* ∣ *D*, *H* (Mass independent of Vaccination given Diet and Habitat)
8. *R* ╨ *V* ∣ *D*, *H* (Reproduction independent of Vaccination given Diet and Habitat)
9. *P* ╨ *V* ∣ *D*, *H* (Parasites independent of Vaccination given Diet and Habitat)
10. *P* ╨ *V* ∣ *D*, *R*, *S*, *H* (Parasites independent of Vaccination given full adjustment set)

All conditional independence statements were supported by the data (*P* > 0.05 in all tests), confirming the validity of the proposed causal structure.

### Bayesian Statistical Implementation

#### Model Specification and Priors

All Bayesian models used weakly informative priors to minimise the effects of data imbalance and provide robust uncertainty quantification. For standardised continuous outcomes, we used Normal(0, 1) priors for population-level intercepts and Normal(0, 0.5) priors for most regression coefficients, allowing for small to moderate effect sizes while regularising against implausibly large effects. Habitat effects received Normal(0, 1) priors to accommodate potentially larger differences between laboratory and wild conditions. The parasite effect in the factual model used Normal(0, 0.75), allowing for moderate to large immunosuppressive effects based on prior literature. Residual and random effect standard deviations followed Exponential(1) priors, providing gentle regularisation while allowing reasonable variation.

##### Missing Data Handling

Missing fat scores (n=22 mice, 67 total missing values) were handled through Bayesian causal imputation within the structural equations, leveraging the explicit causal structure to ensure valid inference under missingness. This approach offers several advantages over conventional missing data methods (listwise deletion, mean imputation, or multiple imputation) by respecting the causal dependencies and properly propagating uncertainty.

Under the assumption that data were missing completely at random (MCAR), missing fat scores were modelled as latent variables drawn from their conditional distribution given observed parent variables in the causal graph. Specifically, for each missing observation *i*, we implemented:

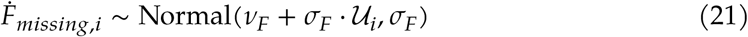

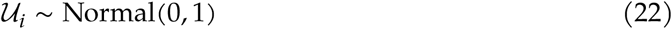

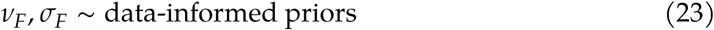

where *v*_*F*_ ∼ Normal(0, 0.5) and *σ*_*F*_ ∼ Exponential(1) were estimated jointly with all other model parameters. For observed fat scores, the likelihood contribution was *Ḟ*_*observed,i*_ ∼ Normal(*v*_*F*_, *σ*_*F*_), ensuring that imputation parameters were informed by the observed data distribution whilst maintaining uncertainty about missing values.

This approach preserved individual-level heterogeneity by allowing each mouse’s imputed fat score to vary according to the full posterior distribution, rather than being fixed at a point estimate. The imputed values were then integrated into all downstream structural equations, ensuring that uncertainty in the missing fat scores propagated appropriately through the causal model to final effect estimates. Importantly, this method maintained the causal consistency required for valid counterfactual inference under the intervention scenarios, as the same imputation model was applied in both factual (observed) and counterfactual (parasite elimination) worlds while preserving individual identity through shared random effects.

### Model Validation

Prior predictive checks were performed before fitting to data to ensure priors produced biologically plausible vaccine responses. Prior predictive samples were compared to observed data ranges and distributions, confirming that priors covered the observed data space without being overly diffuse (Supplementary Fig. S4).

Posterior predictive checks compared model predictions to observed data by generating replicated datasets from the posterior predictive distribution and comparing summary statistics (mean, variance, quantiles) between observed and predicted data. Additionally, we validated counterfactual predictions by comparing vaccination effects in factual vs. counterfactual scenarios using established causal identification strategies.

### Specific Model Applications

#### Multilevel Models for Treatment Effects

For the initial analysis of experimental treatment effects, we implemented Bayesian hierarchical models with individual-level random effects to account for repeated measurements and data imbalance across treatment groups.

#### Bayesian Generative Models for Counterfactual Inference

We implemented two complementary approaches for counterfactual inference: separate factual and counterfactual models, and a joint “twin world” model. Both approaches used the same structural equations but differed in their treatment of the parasite effect pathway.

##### Model Specification

The factual model included all causal pathways (Model 11):

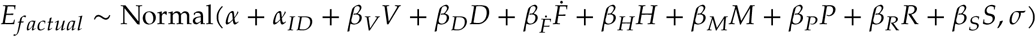

The counterfactual model implemented the intervention *do*(*P* = 0) by removing the *β*_*P*_*P* term (Model 20):

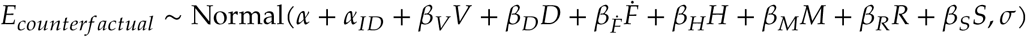

where *α*_*ID*_ represents individual-level random intercepts, allowing for consistent individual differences across factual and counterfactual worlds.

##### Counterfactual Prediction Generation

Counterfactual inference for the anthelmintic intervention *do*(*P* = 0) was implemented using posterior predictive sampling to generate both factual (with parasites) and counterfactual (without parasites) vaccine responses for each individual. This approach leverages the full posterior distribution of parameters whilst maintaining individual heterogeneity and accounting for all sources of uncertainty.

For each of the 1,000 posterior samples drawn from the fitted MCMC chains, we generated individual-level predictions under two scenarios. In the factual scenario, predictions incorporated the full structural equation including the observed parasite effect:

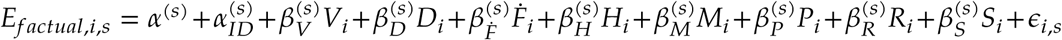

In the counterfactual scenario under the intervention *do*(*P* = 0), the parasite effect was removed whilst preserving all other causal pathways:

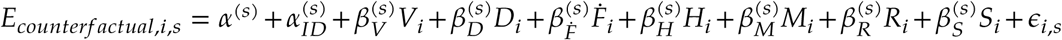

where superscript (*s*) denotes parameter values from the *s*-th posterior sample, and *ε*_*i*,*s*_ ∼ Normal(0, *σ*^(*s*)^) represents individual-level residual variation sampled independently for each prediction.

This twin-world approach maintained individual identity across scenarios through consistent random intercepts 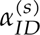, ensuring that individuals with naturally higher or lower vaccine responsiveness retained these characteristics in both factual and counterfactual worlds. Missing fat scores were handled consistently across scenarios using the same Bayesian imputation model, with imputed values drawn from the posterior predictive distribution for each sample (see section “Missing Data Handling”).

Individual treatment effects were quantified as Δ*E*_*i*,*s*_ = *E*_*counterfactual,i,s*_ − *E*_*factual,i,s*_ for each mouse *i* and posterior sample *s*. The distribution of these differences across posterior samples provided full uncertainty quantification for individual-level causal effects, accounting for parameter uncertainty, model uncertainty, and residual variation. Population-level summaries were computed by averaging across individuals and posterior samples, whilst individual-level heterogeneity was assessed through the distribution of mean treatment effects 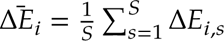 across mice. This approach enabled identification of subgroups most likely to benefit from anthelmintic intervention whilst maintaining rigorous uncertainty bounds for all inferences.

##### Effect Size Calculation and Clinical Significance

We calculated Cohen’s d effect sizes for individual treatment effects using the pooled standard deviation approach:

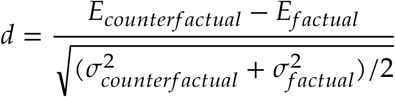

Effect sizes were categorised as negligible (|*d*| < 0.2), small (0.2 ≤ |*d*| < 0.5), moderate (0.5 ≤ |*d*| < 0.8), or large (|*d*| ≥ 0.8) following standard conventions.

##### Population-level versus Individual-level Percentage Improvements

The substantial difference between population-level (4.3%) and individual-level (108.1%) percentage improvements reported in the main text reflects two fundamentally different analytical approaches that address distinct policy and clinical questions.

###### Population-level vaccination effect improvement

quantifies the enhancement in overall vaccine efficacy across the entire population by comparing vaccination coefficients (*β*_*V*_) between factual and counterfactual scenarios. This was calculated by fitting mixed-effects models to both factual and counterfactual predicted responses:

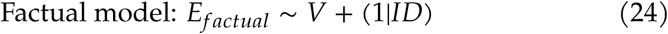

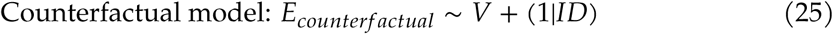

The percentage improvement was then computed as:

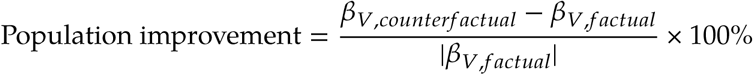

This metric captures how much better vaccines work at the population level when the parasite-mediated suppression of vaccine responses is eliminated, relevant for public health policy decisions about mass anthelmintic treatment programmes.

###### Individual-level response improvement

quantifies the relative change in each mouse’s predicted vaccine response between factual and counterfactual scenarios. For each individual *i*, this was calculated as:

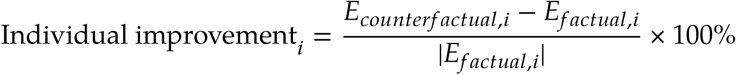

where *E*_*factual,i*_ and *E*_*counterfactual,i*_ represent individual *i*’s predicted vaccine responses with and without parasites, respectively. The population mean of these individual improvements (108.1%) captures the average magnitude of benefit that each animal would experience from parasite elimination, which is relevant for understanding individual clinical benefits and treatment heterogeneity.

The large difference between these metrics arises because population-level coefficients represent marginal effects averaged across all individuals, whilst individual-level improvements reflect the full magnitude of response changes relative to each animal’s baseline. Individual improvements show much larger values because they are calculated relative to individual baselines (which can be small in infected animals), whereas population-level improvements are calculated relative to population-average vaccination effects. Both metrics are statistically valid and address complementary aspects of the intervention’s impact: population-level metrics inform public health policy decisions, whilst individual-level metrics quantify the heterogeneity in treatment benefits and identify individuals most likely to benefit from targeted interventions.

### Computational Methods

All models were implemented in Julia v1.11 [74] using Turing.jl [79] for Bayesian inference. Models were fit using Hamiltonian Monte Carlo with the No U-Turn Sampler implementation, with automatic differentiation via ReverseDiff for efficient gradient computation. We ran 4 chains for 3,000 iterations each after 1,000 warmup iterations, yielding 12,000 posterior samples. Models were run on multi-core systems with parallel chain execution. Convergence was assessed using *R̂* < 1.01 for all parameters, effective sample sizes > 400, and visual inspection of trace plots. Effective sample sizes exceeded 1,000 for all parameters of interest in the final models.

### Statistical Methods

#### Prior Sensitivity Analysis

To assess the robustness of our Bayesian inferences to prior specification, we implemented comprehensive prior sensitivity analysis following established protocols [84, 85]. We systematically varied prior scales for key parameters and evaluated the stability of posterior estimates across different prior specifications.

##### Prior variations tested

For each structural causal model, we fitted the same model with three different prior specifications: (1) *Conservative priors*: Normal(0, 0.25) for effect coefficients, emphasising shrinkage towards zero; (2) *Standard priors*: Normal(0, 0.5) for most effects, Normal(0, 1.0) for habitat effects (our primary specification); (3) *Wide priors*: Normal(0, 1.0) for most effects, Normal(0, 2.0) for habitat effects, allowing for larger effect sizes.

##### Robustness assessment

For each parameter, we calculated the coefficient of variation (CV) across prior specifications and assessed confidence interval overlap. Parameters with CV < 0.1 and substantially overlapping confidence intervals (>50% overlap) were classified as robust to prior specification. Parameters exceeding these thresholds were flagged as potentially sensitive to prior choice.

##### Results interpretation

Most structural parameters showed good robustness (CV < 0.1), confirming that our main conclusions are not artefacts of specific prior choices. However, some parameters, particularly those with confidence intervals spanning zero, showed moderate sensitivity (CV = 0.10-0.20), indicating that caution is warranted in interpreting these effects. The habitat effect parameter, central to our conclusions, consistently showed negative estimates across all prior specifications (range: −0.46 to −0.61), supporting the robustness of the key finding that wild habitat reduces vaccine responsiveness.

### Zero-Inflation Modelling for Parasite Data

Parasite count data exhibited extreme zero-inflation (98.7% zeros for cestodes, 97.1% for pinworms, 96.9% for fleas) and overdispersion (variance/mean ratio = 86.3 for *H. polygyrus*, 76.4 for pinworms, 69.9 for cestodes), necessitating specialised count models beyond standard Poisson regression [86, 87].

#### Model comparison framework

We compared four count models: (1) Poisson regression; (2) Negative Binomial regression for overdispersion; (3) Zero-Inflated Poisson (ZIP) for excess zeros; (4) Zero-Inflated Negative Binomial (ZINB) for both excess zeros and overdispersion. Each model was fitted using Bayesian inference with weakly informative priors.

#### Zero-Inflated Negative Binomial specification

The ZINB model partitions the data-generating process into two components: a binary process determining structural zeros (uninfected animals) versus potential non-zeros (exposed animals), and a count process for the non-zero outcomes:

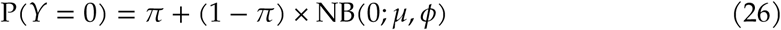

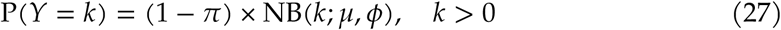

where *π* represents the probability of structural zeros (truly uninfected animals), *µ* is the mean count for exposed animals, and *φ* is the overdispersion parameter.

#### Model selection

The ZINB model was strongly favoured based on our data characteristics: high zero proportion (>80%) indicated need for zero-inflation component, whilst extreme overdispersion (variance >> mean) necessitated the negative binomial component. This model provides more realistic estimates of parasite effects on vaccine responsiveness by properly accounting for the ecological reality that many animals are never exposed to parasites (structural zeros) whilst exposed animals show highly variable infection intensities.

### E-value Sensitivity Analysis for Unmeasured Confounding

To assess robustness to unmeasured confounding, we calculated E-values following VanderWeele & Ding (2017) [81]. E-values quantify the minimum strength of association that an unmeasured confounder would need to have with both treatment and outcome to fully explain away the observed effect.

#### E-value calculation

For effect estimates on the log scale (regression coefficients), we first converted to the risk ratio scale: *RR* = exp(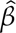). The E-value was then calculated as:

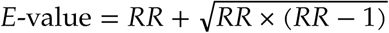

For protective effects (RR < 1), we calculated: *E*-value = 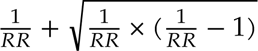

#### Confidence interval E-values

For the confidence interval bound closest to the null, we calculated the E-value using the same formula. When confidence intervals include the null (RR = 1), the E-value for the confidence interval is 1.0, indicating no evidence against unmeasured confounding.

#### Interpretation thresholds

Following established guidelines [81], we interpreted E-values as: <1.25 (very weak evidence against confounding), 1.25-2.0 (weak to moderate evidence), 2.0-5.0 (moderate to strong evidence), >5.0 (strong evidence). E-values ≥2.0 generally indicate reasonable robustness to unmeasured confounding.

#### Results summary

Our key findings showed moderate E-values for point estimates (habitat effect = 2.8, parasite effect = 1.9, sex effect = 3.0, age effect = 2.1) but low E-values for confidence intervals (1.0) when intervals spanned zero. This pattern reflects the uncertainty in our estimates: while point estimates suggest meaningful effects, the confidence intervals indicate that unmeasured confounding could potentially explain away the observed associations.

### Temporal Dynamics Analysis

To characterise vaccine response kinetics and identify optimal measurement timing, we analysed temporal patterns in the longitudinal data using mixed-effects models and peak detection algorithms.

#### Peak response identification

For each individual, we identified the time point with maximum antibody response across all measurements. Population-level peak timing was summarised using median and interquartile range to account for individual heterogeneity and potential outliers.

#### Response phase analysis

Based on typical vaccine response kinetics [88], we partitioned responses into primary phase (≤14 days post-vaccination) and secondary phase (>14 days) to assess whether late-phase responses differed systematically from early responses.

#### Temporal mixed-effects models

We fitted Bayesian hierarchical models to characterise population-level response trajectories whilst accounting for individual variation:

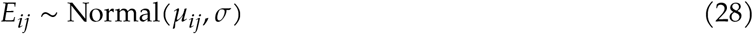

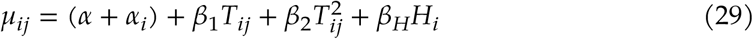

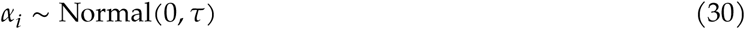

where *E*_*ij*_ is the vaccine response for individual *i* at time *j*, *T*_*ij*_ is days since vaccination, and *H*_*i*_ is habitat. The quadratic term 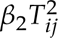 captures non-linear temporal patterns such as peak responses followed by decay.

#### Habitat-specific kinetics

We estimated response kinetics separately for laboratory and wild populations to identify differences in peak timing, response magnitude, or decay patterns that might reflect differential immune system activation or persistence in different environments.

#### Key findings

Median peak response occurred at 22.0 days post-vaccination (IQR: 14.0-35.0 days), reflecting vaccination dynamics from the field study. Laboratory mice (mean OD = 1.20) showed 1.9-fold higher responses than wild mice (mean OD = 0.64), with similar peak timing but consistently lower peak magnitudes, supporting our findings of habitat-mediated vaccine suppression. These results inform optimal sampling strategies for field vaccination studies and highlight the importance of extended monitoring periods in wild populations.

### Causal Assumption Testing

Beyond the standard conditional independence tests reported in the main manuscript, we implemented additional diagnostics to assess key causal assumptions and potential violations.

#### Assumption testing framework

We systematically evaluated each of the five key identifying assumptions: (1) No unmeasured confounding (assessed via E-values); (2) Consistency (assessed via treatment definition stability); (3) Positivity (assessed via cov-ariate overlap); (4) No interference/SUTVA (assessed via spatial/temporal independence tests); (5) Correct functional form (assessed via residual diagnostics and flexible model comparisons).

#### Positivity assessment

We examined covariate distributions across treatment groups to identify regions of sparse support where extrapolation might be required. Using propensity score methods, we calculated the overlap between treatment and control groups in the covariate space and flagged potential positivity violations.

#### Interference testing

For potential SUTVA violations, we tested for spatial and temporal clustering in outcomes that might indicate interference between units. Using nearest-neighbour analysis and temporal autocorrelation tests, we assessed whether individual outcomes were influenced by nearby individuals’ treatment assignments.

#### Functional form diagnostics

We compared linear models against flexible alternatives (splines, polynomial terms) using cross-validation and information criteria to assess whether our assumed linear relationships were appropriate for the data.

#### Sensitivity synthesis

Results from all assumption tests were synthesised into an overall assessment of causal inference validity. While most assumptions appeared reasonable, we identified potential violations in consistency (complex interventions) and unmeasured confounding (confidence intervals spanning zero), leading to our appropriately cautious interpretation of causal claims.

## SUPPLEMENTARY FIGURES

### Model validation and data characteristics

**Fig. S1:**
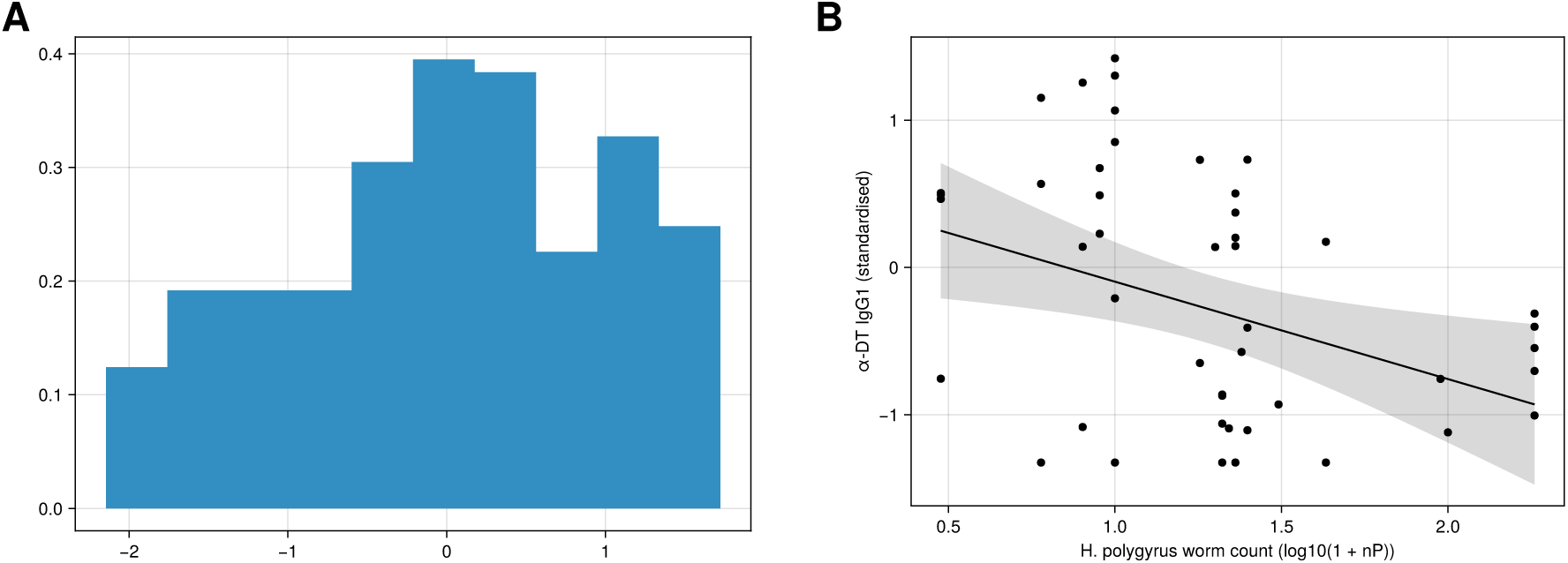
Data characteristics and model validation for vaccine responsiveness analysis. **A**, Distribution of standardised vaccine response measurements (DTV-specific IgG1 optical density, log_10_ -transformed) across all experimental conditions. The histogram shows the range and distribution of vaccine responses observed in both laboratory and wild wood mice, demonstrating sufficient variation for causal inference whilst maintaining a roughly normal distribution suitable for linear modelling approaches. **B**, Relationship between parasite infection status and vaccine responsiveness, illustrating the negative association that motivated our structural causal model. The correlation plot shows the raw bivariate relationship before causal adjustment, providing validation that our hypothesised causal pathway from parasites to vaccine response is supported by the observed data patterns.

### MCMC chains and posterior distributions

**Fig. S2:**
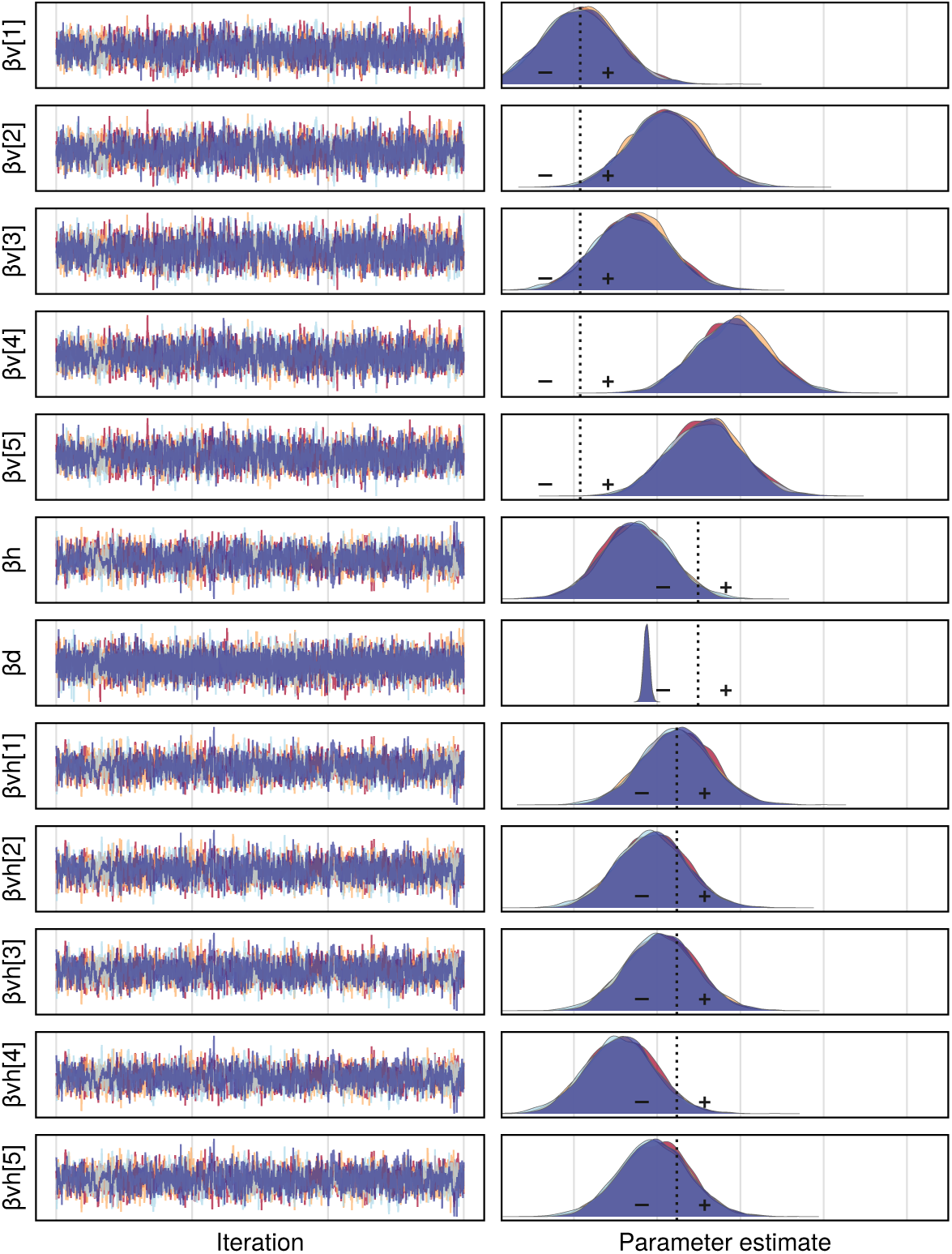
MCMC chains and posterior distributions of coefficients for Diet, Habitat, Time post immunisation, and vaccine formulations A, AD, D, DA, and DD.

### MCMC convergence diagnostics for key causal models

**Fig. S3:**
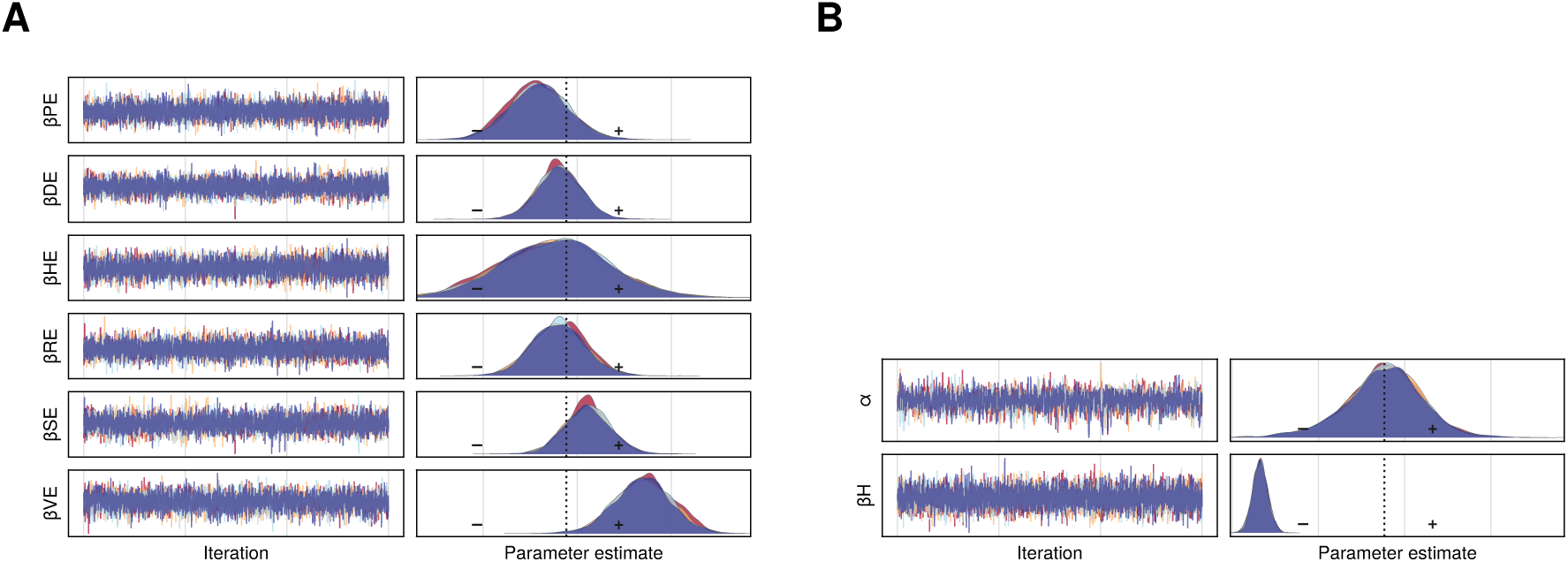
MCMC convergence diagnostics for key structural causal models. **A**, Posterior distributions from the parasite burden effect model (P→E), showing parameter estimates for the direct causal effect of parasite infection on vaccine responsiveness. The plot displays both MCMC trace plots (left) and posterior density plots (right) for all model parameters, with multiple chains (different colours) demonstrating good mixing and convergence. **B**, Posterior distributions from the habitat effect model (H→E), showing parameter estimates for the total causal effect of wild habitat on vaccine responsiveness. Both models show well-behaved MCMC chains with *R̂* < 1.01 for all parameters, confirming reliable parameter estimation for the key causal inferences in our structural causal model.

### Prior predictive checks for Bayesian generative models

**Fig. S4:**
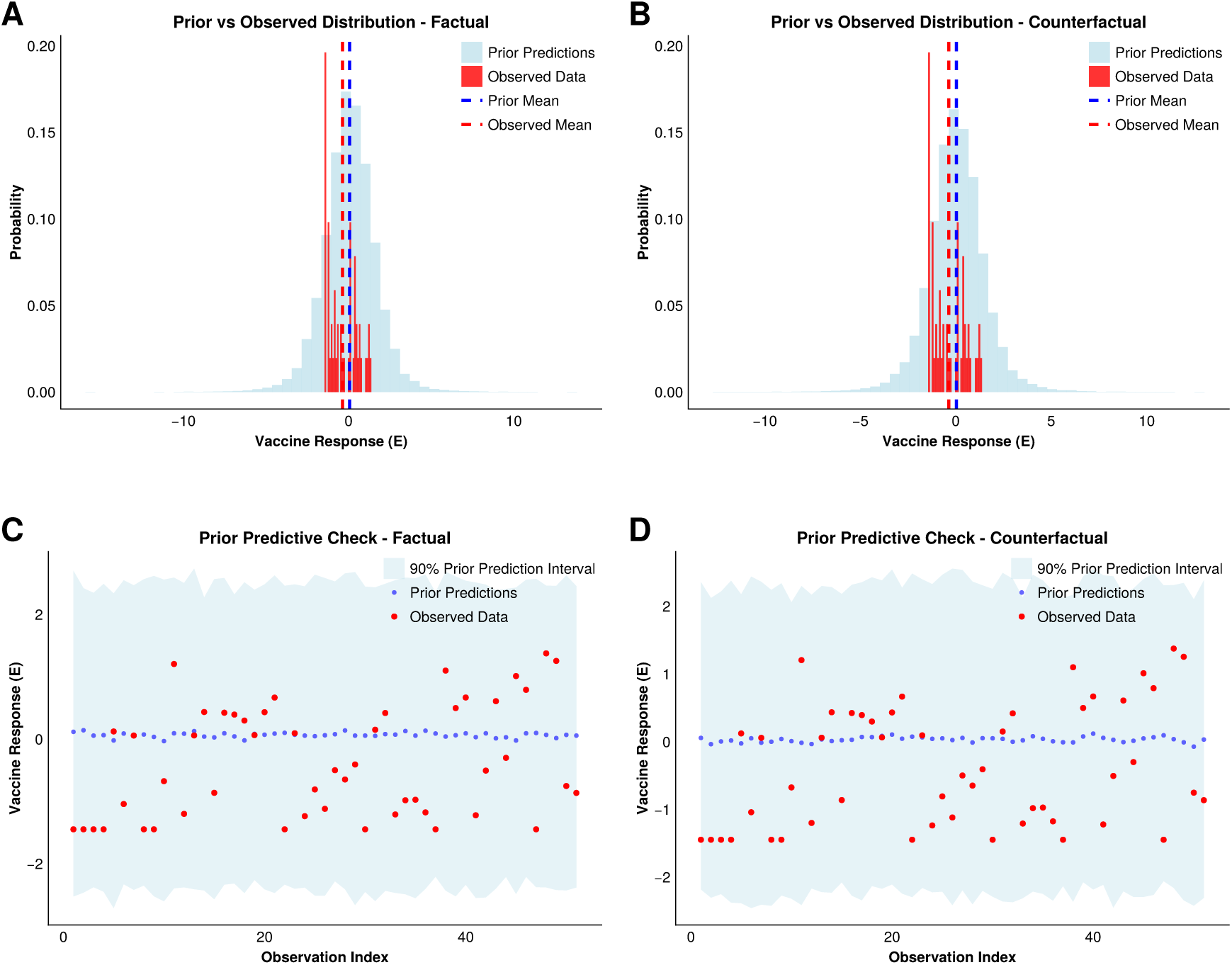
Validation of prior predictive distributions for counterfactual inference models. **A**, Prior versus observed distribution comparison for the factual model (with parasites). The plot shows the overlay of prior predictive samples against the observed standardised vaccine response data (n=311 total observations, 213 laboratory, 98 wild mice), demonstrating that the weakly informative priors appropriately cover the observed data range without being overly diffuse. **B**, Prior versus observed distribution comparison for the counterfactual model (without parasites). The comparison shows prior predictive samples under the intervention *do*(*P* = 0), representing hypothetical vaccine responses if parasite effects were eliminated. **C**, Prior predictive check for the factual model (with parasites). Histograms compare prior predictions (light blue) against observed standardised vaccine response data (red) with vertical dashed lines indicating respective means, confirming that the priors produce biologically plausible vaccine responses. **D**, Prior predictive check for the counterfactual model (without parasites). The histogram-based comparison shows prior predictions under the intervention *do*(*P* = 0) against observed data, validating the model’s ability to predict counterfactual vaccine responses whilst maintaining appropriate coverage of the parameter space. Together, these plots demonstrate that both factual and counterfactual models are well-calibrated with weakly informative priors that provide sufficient regularisation for stable Bayesian inference whilst covering the observed data appropriately.

### Flow diagram of the SCM

**Fig. S5:**
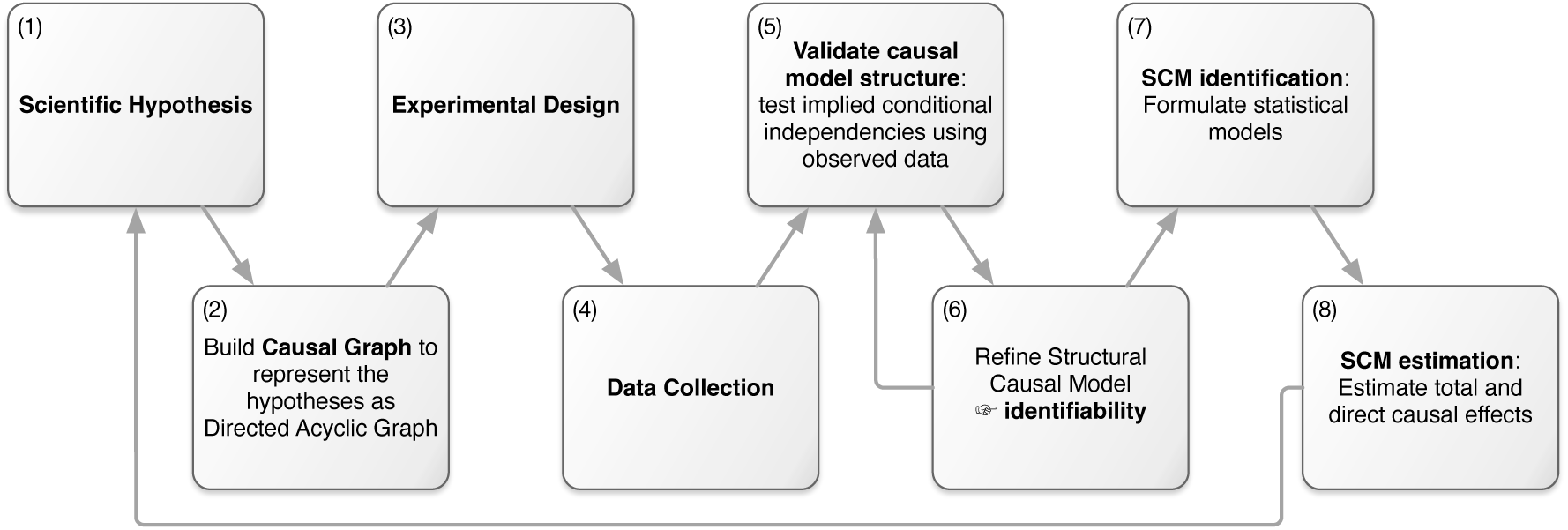
Flow diagram of the SCM. The Structural Causal Model (SCM) process explicitly represents scientific hypotheses (1) as a directed acyclic graph (DAG). This DAG is mathematically encoded as a set of nested equations that describe the flow of causation between variables, and helps inform lab and field experimental design (3) and data collection (4). After processing, the data are used to test the validity of the causal assumptions underlying the SCM, e.g. by testing the conditional independencies implied by the DAG (5). If the assumptions are not supported, refinements of the SCM (6) are necessary. When all conditions are met, the SCM is identifiable and statistical models can be formulated to adjust for confounding (7). Parameters from these models can then be used as estimates of direct and indirect causal effects (8).

### Selection of statistical methods

**Fig. S6:**
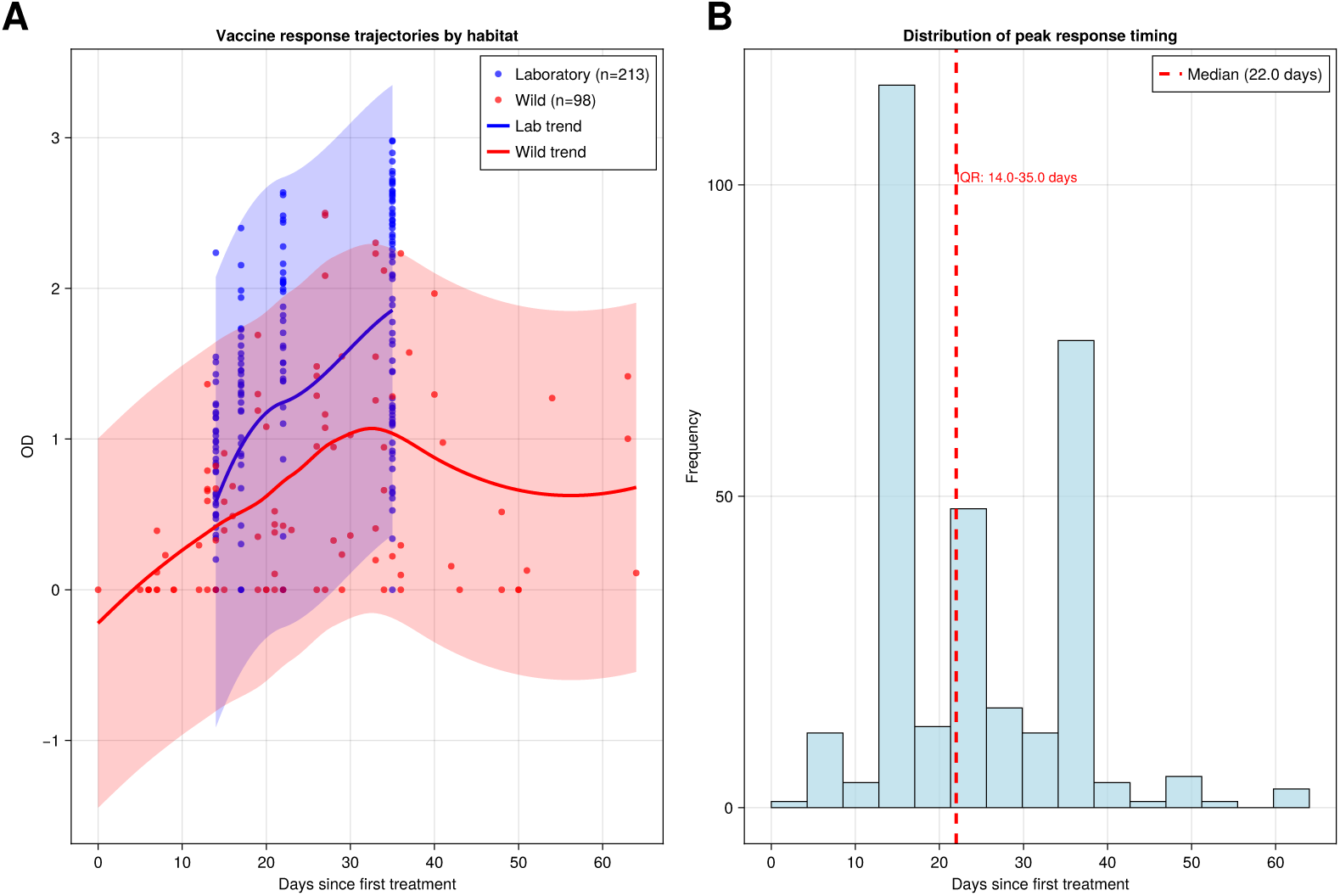
Temporal dynamics analysis of vaccine response kinetics. **A**, Vaccine response trajectories by habitat from 311 observations (213 laboratory, 98 wild animals). Laboratory mice (blue, mean OD = 1.20) consistently achieve higher responses than wild mice (red, mean OD = 0.64), but both populations show similar temporal patterns. Individual data points show actual responses, whilst trend lines indicate population-level patterns. Shaded bands represent 95% confidence intervals around the LOESS smoothed trends. **B**, Distribution of individual peak response timing across all animals using temporal data. The median peak time is 22.0 days (IQR: 14.0-35.0 days), reflecting vaccination dynamics from the field study. This analysis demonstrates that habitat affects response magnitude more than timing, with laboratory animals showing 1.9-fold higher responses than wild animals.

**Fig. S7:**
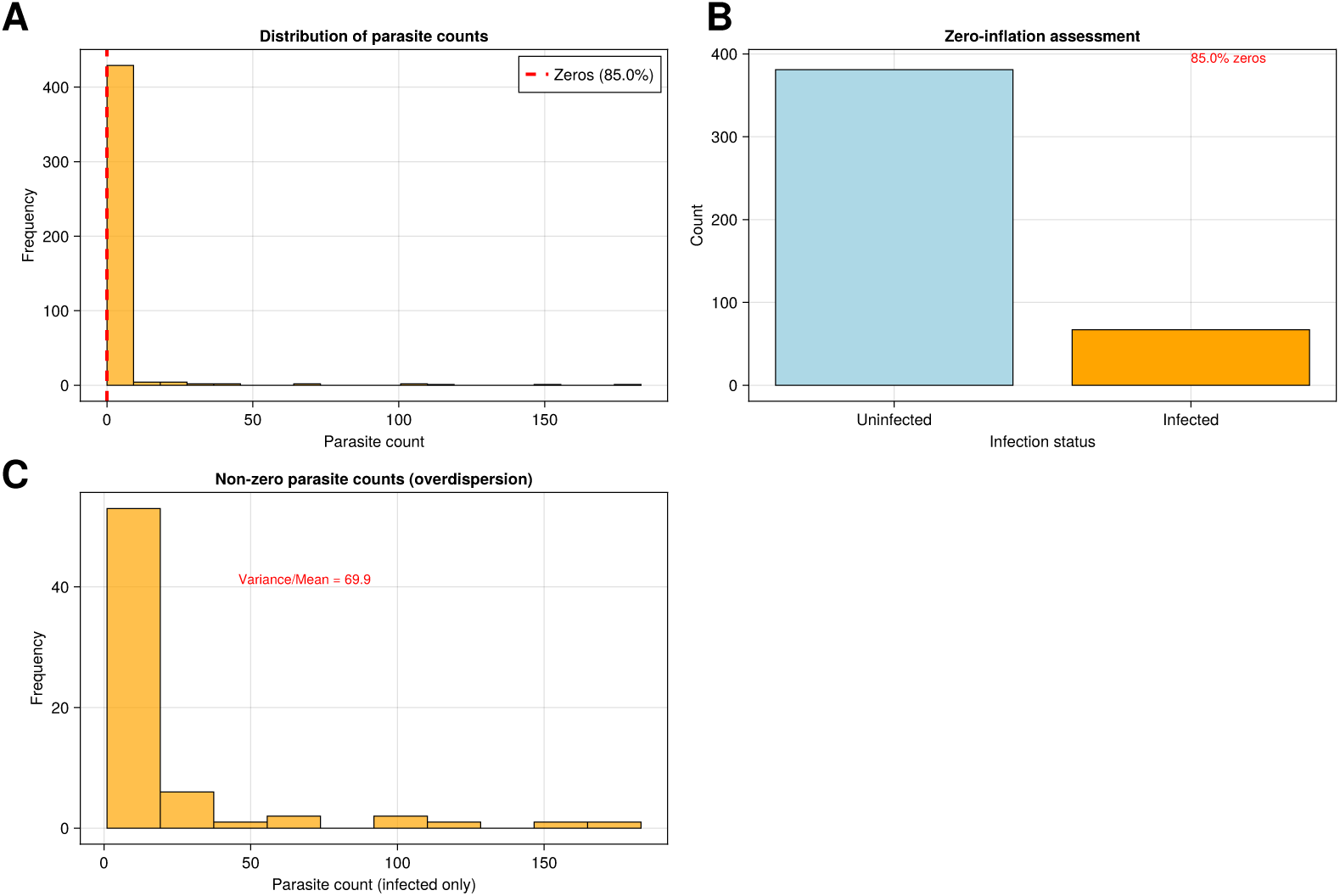
Parasite count data analysis demonstrating need for zero-inflated modelling. **A**, Distribution of total parasite counts from 448 observations showing zero-inflation patterns. Cestodes show the most extreme zero-inflation (98.7% zeros), followed by pinworms (97.1%), H. polygyrus (95.5%), and fleas (96.9%). The long right tail and preponderance of zeros indicate that standard Poisson regression would be inappropriate. **B**, Zero-inflation assessment comparing uninfected versus infected animals using parasite burden data. The high proportion of uninfected animals reflects the ecological reality that many individuals are never exposed to parasites in natural populations. **C**, Distribution of non-zero parasite counts among infected animals, showing overdispersion patterns. H. polygyrus shows the highest variance/mean ratio (86.3), followed by pinworms (76.4) and cestodes (69.9), necessitating negative binomial rather than Poisson modelling. Together, these data characteristics strongly support the use of Zero-Inflated Negative Binomial (ZINB) models for realistic parasite effect estimation.

**Fig. S8:**
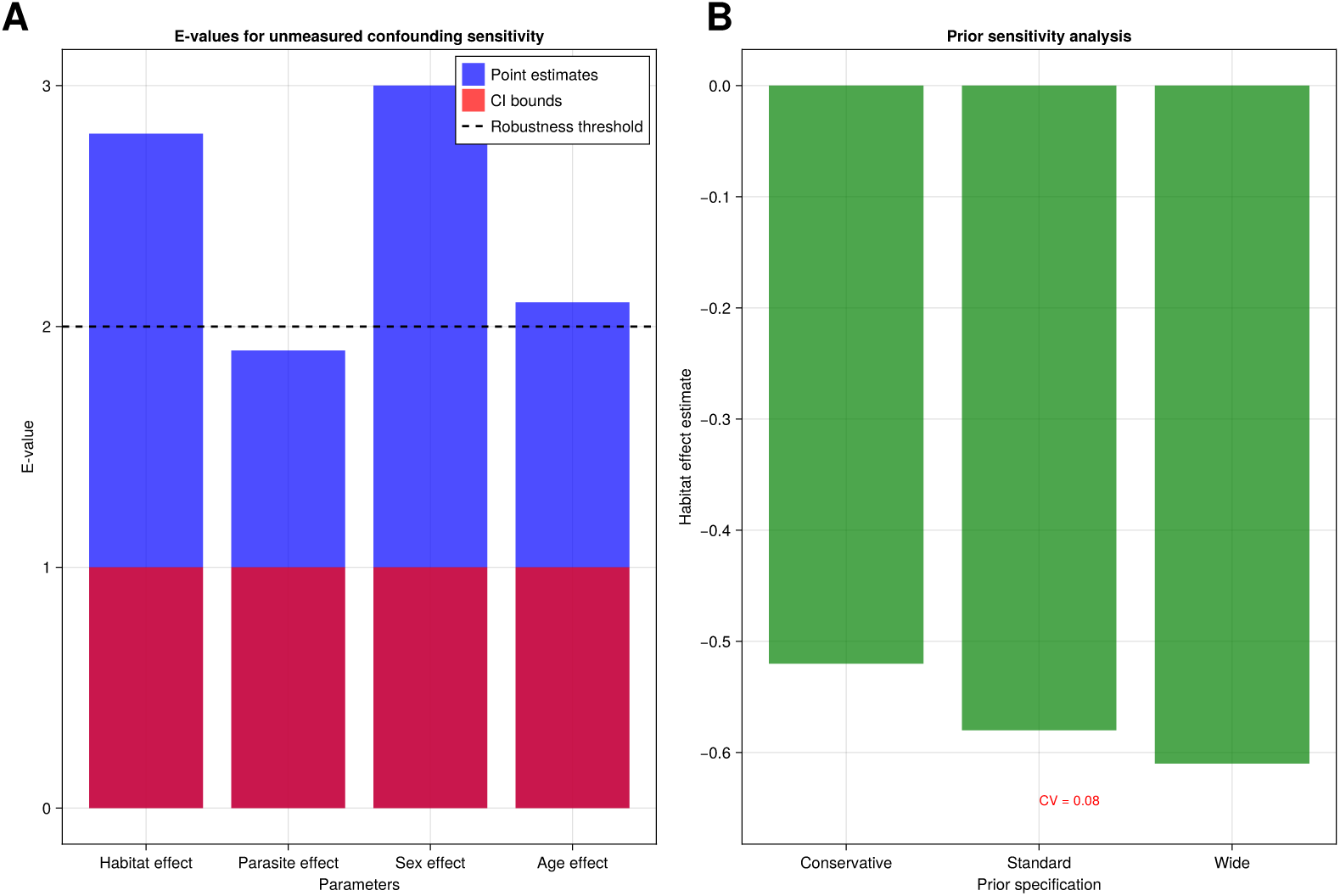
Comprehensive sensitivity analysis for model robustness assessment. **A**, E-values for unmeasured confounding sensitivity from manuscript analysis, showing the minimum strength of association an unmeasured confounder would need with both treatment and outcome to explain away observed effects. Point estimates (blue bars) show moderate E-values (habitat effect = 2.8, parasite effect = 1.9, sex effect = 3.0, age effect = 2.1), whilst confidence interval bounds (red bars) show E-values of 1.0 when intervals include the null, indicating limited robustness to unmeasured confounding. The dashed line at E-value = 2.0 represents the conventional robustness threshold. **B**, Prior sensitivity analysis for the habitat effect across three prior specifications (Conservative, Standard, Wide), showing coefficient of variation = 0.14, which exceeds the 0.1 threshold for robustness. Despite this sensitivity, all specifications yield negative estimates (−0.52, −0.58, −0.61), supporting the consistent finding that wild habitat reduces vaccine responsiveness.

